# A proactive genotype-to-patient-phenotype map for cystathionine beta-synthase

**DOI:** 10.1101/473983

**Authors:** Song Sun, Jochen Weile, Marta Verby, Atina G. Cote, Yingzhou Wu, Iosifina Fotiadou, Julia Kitaygorodsky, Jasper Rine, Pavel Ješina, Viktor Kožich, Frederick P. Roth

## Abstract

Success in precision medicine depends on our ability to determine which rare human genetic variants have functional effects. Classical homocystinuria—characterized by elevated homocyst(e)ine in plasma and urine—is caused by primarily-rare variants in the cystathionine beta-synthase (*CBS*) gene. About half of patients respond to vitamin B_6_ therapy. With early detection in newborns, existing therapies are highly effective. Functional *CBS* variants, especially those that respond to vitamin B_6_, can be detected based on their ability to restore growth in yeast cells lacking *CYS4 (*the yeast ortholog of *CBS*). This assay has previously been carried out only ‘reactively’ after first observation of a variant in patients. Here we describe a ‘proactive’ comprehensive missense variant effect map for human CBS. Together, saturation codon-replacement mutagenesis, *en masse* growth selection at different vitamin B_6_ levels, and sequencing yielded a ‘look-up table’ for CBS missense variant function and vitamin B_6_-remediability in yeast. The CBS variant effect map identified disease variants and predicted both disease severity (*r* = 0.82) and human clinical response to vitamin B_6_ (*r* = 0.89). Thus, highly-multiplexed cell-based assays can yield proactive maps of variant function and patient response to therapy, even for rare variants not previously seen in the clinic.

## Introduction

Rapid development of high-throughput sequencing technology has made it feasible to sequence the genome of every human. However, pressing challenges remain in interpreting the clinical impact of genetic variants discovered by sequencing and ultimately in personalizing diagnostic surveillance and therapy. Over 138,000 exomes have been collected in the Genome Aggregation Database (gnomAD)^1^ and 4.6 million coding variants have been discovered. Among these discovered coding variants, 99% are rare, with a minor allele frequency (MAF) below 0.5%. Although statistical association methods have identified many common variants that correlate with (and in some cases cause) human disease, correlational methods are typically futile for rare variants. In ClinVar^2^, only 2% of rare coding variants have a clinical interpretation and the majority of interpreted variants were annotated as “variants of uncertain significance” (VUS)^3^.

Diverse computational and experimental methods have been developed to predict the functional impact of rare coding variants. Many computational methods can score all possible missense variants ‘proactively’ and thus make supporting evidence for variant interpretation available immediately upon variant discovery. However, computational predictions were found to identify fewer than 20% of pathogenic variants when used at stringent thresholds where >90% of pathogenic variant predictions were correct^4^. At permissive thresholds that detect 90% of pathogenic variants, fully ∼30% of disease variant predictions were erroneous^4^. More accurate predictions can come from experimentally interrogating the functionality of each variant^4^, but this one-at-a-time approach is prohibitively laborious and time-consuming. Even where done, these experimental assays have necessarily been ‘reactive’, with results that lag far behind first clinical presentation.

Variant effect (VE) mapping^5,6^ is a strategy for testing the function of a large number of variants in a single experiment. A VE map provides a ‘look-up table’ for functionality of coding variants in disease-associated genes, potentially providing strong evidence for immediate clinical interpretation of patient variants^7,8^, meeting a clinical need that is particularly acute for rare and personal variants found via genome sequencing. Although experimental VE maps generally contain some missing data, a recently-published VE-mapping framework used machine learning to impute missing data so that, given a critical mass of experimental data, missing values could be filled in with accuracy approaching that of experimental measurement^9^.

Human cystathionine β-synthase (CBS) is an enzyme that catalyzes the first step in the transsulfuration pathway—condensation of serine and homocysteine to give cystathionine^10^. This reaction eliminates the toxic metabolite homocysteine and, by using the alternative substrate cysteine, also produces hydrogen sulfide, a gaseous signaling molecule^11,12^. CBS forms tetramers and uses both heme and pyridoxal 5′-phosphate (PLP; the active form of vitamin B_6_) as cofactors, and S-adenosylmethionine (AdoMet) as an allosteric activator^13^.

Classical homocystinuria (OMIM # 236200) is an autosomal recessive disorder of methionine metabolism manifested by abnormal accumulation of homocysteine in blood and increased excretion of homocystine in urine, and by simultaneous decrease of plasma cystathionine. The disease was discovered in 1962^14^ and two years later shown to be caused by deficient CBS activity in the liver^15^. Since the identification of the first disease-causing CBS variants^16^, several hundred alleles have been identified from homocystinuria patients^17^, many of which have been further genetically and biochemically characterized^18-23^, yielding ∼200 annotated pathogenic variants^24,25^.

Two major forms of this disease have been described. About half of patients suffers from a severe CBS deficiency which manifests in childhood by lens dislocation (luxation), skeletal abnormalities resembling Marfan syndrome, thromboembolism and neuropsychiatric problems. This type of disease usually does not respond to vitamin B_6_ treatment; however, early initiation of therapy with low methionine diet and/or betaine in newborn period prevents most of the clinical complications ^26^. The other half of patients suffers from the milder form of disease, which typically manifests by thromboembolism in adulthood and which responds to vitamin B_6_ treatment^26-28^.

The population frequency of severe early-onset CBS deficiency ranges from 1 in 60,000 to 1 in 900,000 between countries, the worldwide birth frequency of clinically-ascertained patients was estimated to be 1:122,000^29^. However, homocystinuria may be more frequent in specific populations (1:1,800 in Qatar) and higher frequency of the adult B_6_-responsive form has been also suggested based on molecular epidemiological studies ^30-32^ ^27,28^.

Since only early diagnosis and timely therapy can effectively prevent long-term complications in patients with homocystinuria, many newborn screening programs worldwide target CBS deficiency^33^. Screening by determining total homocysteine (tHcy) in dry blood spots is not routinely done given the need of LC-MS/MS assay and associated costs. Thus, CBS deficiency is usually screened for by detecting elevated methionine concentration, which should be complemented with second tier testing for tHcy^34^. Unfortunately, methionine screening may miss some vitamin B_6_ non-responsive patients, and a large proportion of vitamin B_6_-responsive patients. It is at present unknown whether future newborns screening programs based possibly on genome sequencing will improve the detection of homocystinuria.

Yeast complementation assays can identify pathogenic alleles with high accuracy^4^. The human *CBS* gene can complement growth defects in *cys4*Δ yeast deletion mutants^35,36^, and this assay can also be used to test whether variants are vitamin B_6_-dependent^37,38,39,40^. Here we adapt this complementation assay to our recently-described VE mapping framework, and use it to generate comprehensive functional maps of *CBS* missense variation with low or high levels of vitamin B_6_. We find that scores from the resulting VE maps could identify functional variation in *CBS* and, in an independent patient cohort, patient CBS enzyme activities inferred from the VE map correlated strongly with age of onset, disease severity, and response of CBS deficient patients to vitamin B_6_ therapy.

## Results

To enable interpretation of genetic variation in *CBS*, we sought to test all possible missense variants of *CBS* for functional effects and vitamin B_6_-remediability. We therefore reimplemented a previously validated humanized yeast model^35-39^, confirming that expression of human *CBS* from the hORFeome collection restores the ability of a yeast *cys4*Δ strain to grow without supplementation of glutathione (which provides a source for cysteine that circumvents the need for cystathionine; see Supplementary Fig. 1). Coupling this functional complementation with our recently developed framework for exhaustively mapping functional coding variants, we attempted to test the functional impact as well as the vitamin B_6_ remediability of all possible missense *CBS* variants in parallel. (The overall scheme is described in Fig. 1b).

**Figure 1.**
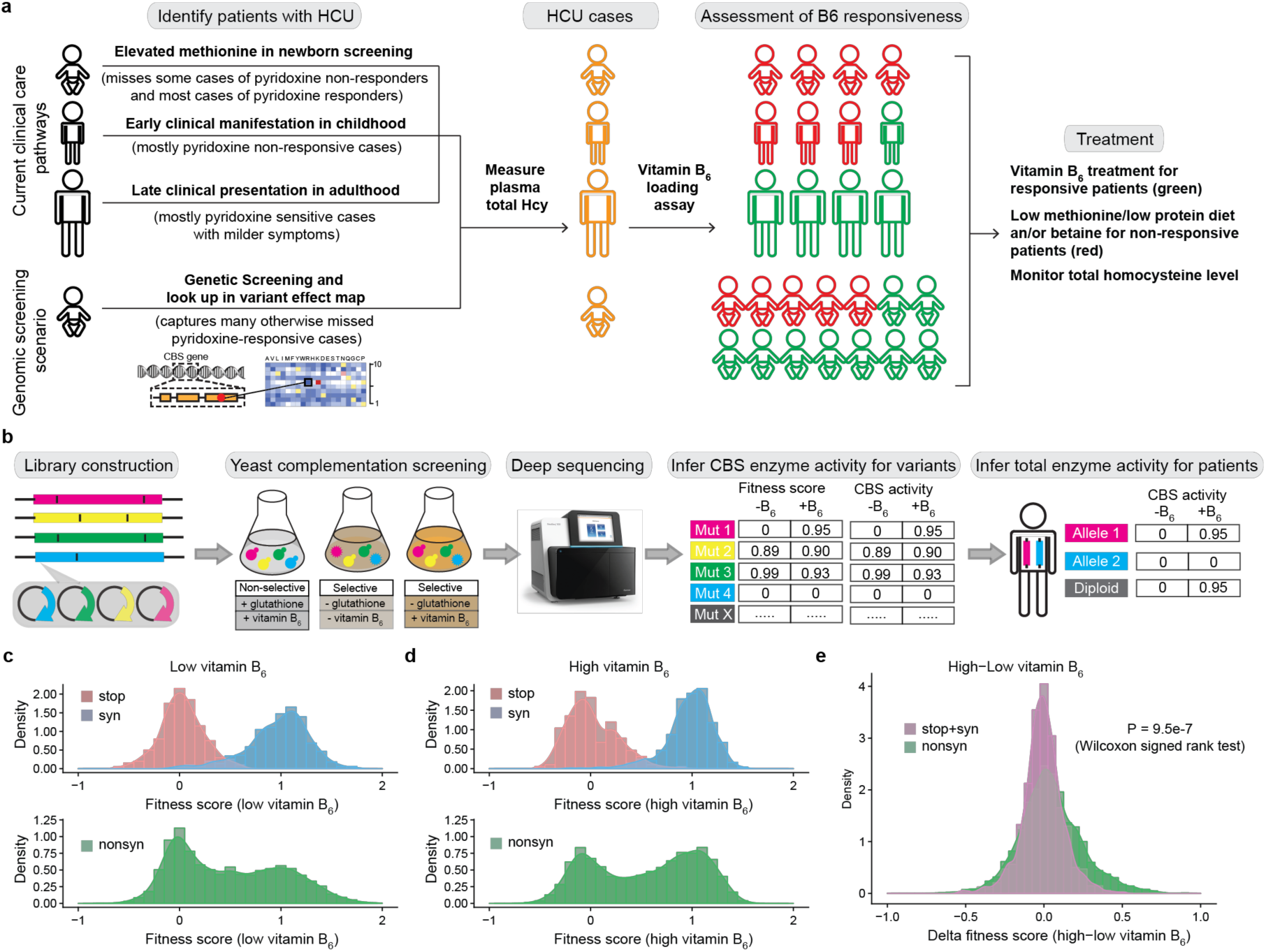
Diagnosis and treatment of homocystinuria (HCU) patients, and production and potential utility of a CBS variant effect map. (a) Current clinical care pathways for identifying, confirming and treating homocystinuria patients (which probably miss many cases of CBS deficiency and especially those that are most responsive to vitamin B_6_ therapy) and a proposed genomic screening scenario. (b) Workflow for generating the CBS variant effect maps using low or high levels of vitamin B_6_ and inferring total enzyme activities for patients. (c-d) Fitness score distributions of stop codon, synonymous and missense variants with low (c) or high (d) levels of vitamin B_6_. (e) Comparison of the distribution of delta scores for missense variants with the null distribution (delta scores for nonsense and synonymous variants).

First, we constructed a library of *CBS* variants using a previously described codon replacement mutagenesis method^9^. The variant library, initially generated as a pool of amplicons, was transferred *en masse* into the appropriate yeast expression vector via two steps of recombinational subcloning. The resulting library of variant expression clones was then transformed *en masse* into the yeast *cys4* mutant strain.

Next, pools of transformed yeast *cys4* mutant strains were grown competitively in cysteine-lacking medium supplemented with low (0 and 1ng/ml) or high (400 ng/ml) concentrations of vitamin B_6_. Allele frequencies of *CBS* variants before and after selection were determined by next-generation sequencing. We used the TileSeq approach^9^, sequencing a tiling set of ∼100 nucleotide segments amplified from the pool. We sought to minimize base-calling error (which can complicate quantitation of low allele frequency variants within a pool) by sequencing both forward and reverse strands of each template cluster on the flow cell and only accepting variants for which the complementary variant on the opposite strand is also seen. Sequencing was performed such that both forward and reverse strands of each nucleotide position were covered by ∼2M reads. In the pre-selection pool, this sequencing detected 83% of all possible missense variants and 94% of the amino-acid substitutions that can be achieved via a single-nucleotide variant (SNV). Fitness scores were calculated for each amino acid substitution based on post-selection changes in allele frequency under both low or high vitamin B_6_ conditions (Materials and Methods), yielding an initial VE map for CBS. To consider only fitness scores where allele frequencies were high enough to be accurately measured, we kept only the 50% of amino acid substitutions with a pre-selection allele frequency above 0.005% (Materials and Methods).

Fitness scores from the resulting VE maps were strongly correlated with the relative growth rates previously determined in single-variant growth assays^39^ with Pearson correlation coefficient (PCC) values over 0.8 (Supplementary Fig. 2a-c). Our results also showed weaker but still-significant correlation with another single variant analysis^41^ (Supplementary Fig. 2d). Because fitness scores were highly correlated between the two screens with low levels of vitamin B_6_ (0 and 1 ng/ml), we combined these two datasets to generate a single set of “low vitamin B_6_” fitness scores (Supplementary Fig. 2e). We also estimated the standard error of each score, and filtered each map further to consider only scores with an estimated standard error below 0.2. After filtering, 49% of all possible missense variants and 76% of all SNV-accessible amino acid substitutions were well measured in the low vitamin B_6_ map. Similarly, 48% of all missense variants and 72% of SNV-accessible substitutions were well measured in the high vitamin B_6_ map.

**Figure 2.**
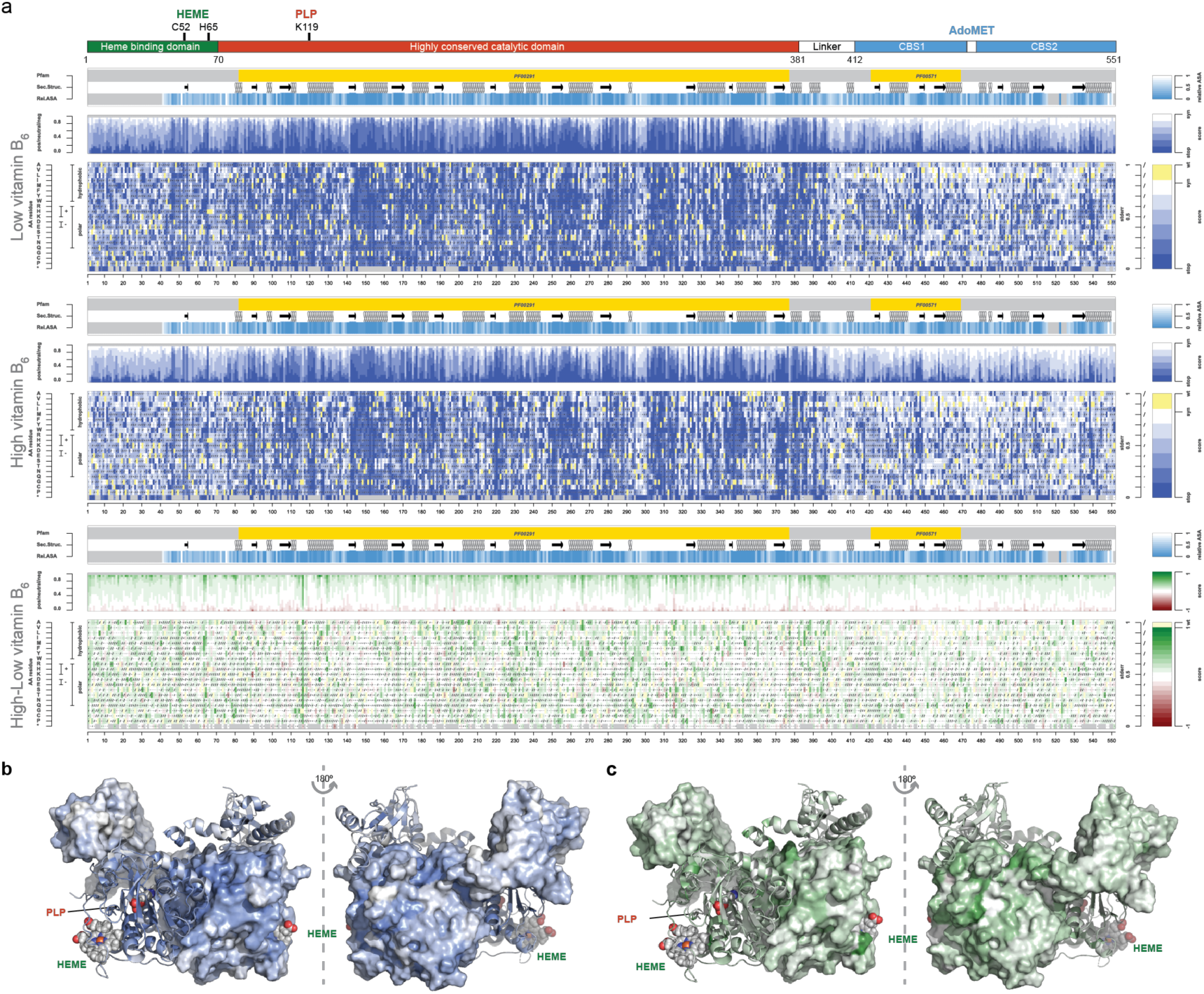
CBS variant effect (VE) maps and accordingly colorized structures of CBS. (a) Complete VE maps for CBS: fitness landscape with low level (top) and (middle) levels of vitamin B_6_, and the delta fitness (high-low vitamin B_6_) landscape (bottom). For high and low vitamin B_6_ VE maps, a functional score of 0 (blue) corresponds to the median fitness of nonsense variants. A score of 1 (white) corresponds to the median fitness of synonymous variants. For the delta fitness landscape (high-low vitamin B_6_), substitutions were colored green if delta fitness score is positive and red if negative. (b-c) Crystal structure of a CBS dimer with residues colored according to the median variant fitness with low vitamin B_6_ (b) or the median delta fitness score (c). The CBS dimer structure shown is based on PDB entry 4L3V^52^ with one monomer in cartoon style and the other monomer in surface mode.

Synonymous variants and nonsense variants each exhibited unimodal fitness score distributions that were well separated from one another (Fig. 1c-d). Missense variants under both selection conditions showed bimodal distributions (Fig. 1c-d). We also calculated a “delta” fitness score (high vitamin B_6_ – low vitamin B_6_ fitness score) for each variant. The distribution of delta fitness scores for missense variants had a longer positive tail than did nonsense and synonymous variants, indicating that the fitness of some missense variants was substantially increased by elevated levels of vitamin B_6_ (Fig. 1e).

Given a critical mass of experimental variant-effect measurements, missing data can be imputed using a machine learning model with confidence that approaches that of experimental measurement^9^ (unpublished result). Therefore, we used a gradient boost tree regression model (unpublished result) to impute missing entries. This yielded complete variant effect maps for CBS under both low and high vitamin B_6_ conditions, which in turn enabled a map of functional remediability of missense variation to different vitamin B_6_ levels (Materials and Methods; Fig. 2a; Supplementary Fig. 3a; Supplementary Dataset 1).

**Figure 3.**
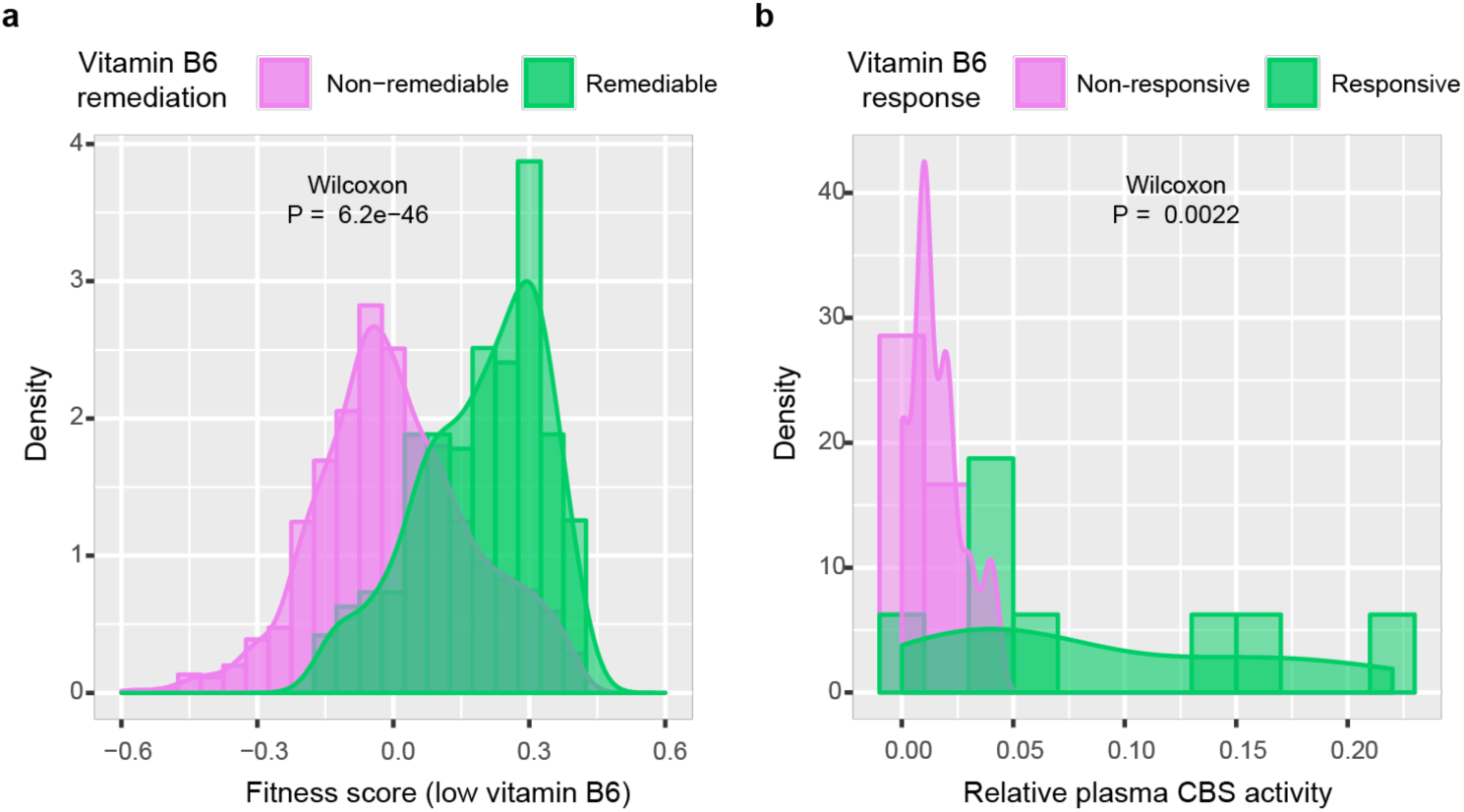
Variant effect maps confirm that vitamin B_6_ is more likely to remediate the weakest-effect variants. (a) Distributions of low vitamin B_6_ fitness scores for variants that were deleterious under the low vitamin B_6_ condition, separated according to whether not they were vitamin B_6_ remediable. (b) Relative plasma CBS activity distributions of vitamin B6-responsive and nonresponsive homocystinuria patients.

The set of CBS variant effect maps were largely consistent with known biochemical and structural features of the CBS protein. Early truncating stop codon variants are uniformly deleterious throughout the whole protein except the small linker region between the catalytic domain and the C-terminal regulatory domain. Concordant with the previous finding that truncating variants at amino acid positions 409 and 410, that are pathogenic in humans, increase CBS basal enzyme activity upon expression in yeast^38^, nonsense variants at these positions exhibited hyper-complementation in both variant effect maps (Supplementary Fig. 3a). Colouring each residue in CBS crystal structure with the median variant fitness at that position shows that residues in the central PLP-binding catalytic domain, and especially those nearest to bound PLP, are intolerant to variation (Fig. 2b). Positions in the heme-binding domain are more tolerant to variation compared to the PLP-binding domain, however, substitutions in the two heme-binding residues Cys52 and His65 are detrimental (Supplementary Fig. 3a; Fig. 2a). The C-terminal AdoMet-activated repressive domain is more tolerant to variation (Fig. 2a-b) suggesting that, at least for the yeast strain and growth media conditions we used, functionality of this domain is not required for yeast complementation.

The ‘delta’ map, measuring high vitamin B_6_ - low vitamin B_6_ fitness, showed that a substantial fraction of missense variants have increased activity at an elevated vitamin B_6_ level. To better understand mechanisms of vitamin B_6_ remediation, we examined the low vitamin B_6_ map to identify variants with fitness scores that were significantly worse than the fitness distribution of synonymous variants (Materials and Methods; Supplementary Fig. 4a). Variants that were deleterious under low vitamin B_6_ conditions were then classified as vitamin B_6_-remediable or non-remediable according to whether their delta fitness score significantly deviated from the distribution of delta scores for nonsense variants (Materials and Methods; Supplementary Fig. 4b).

**Figure 4.**
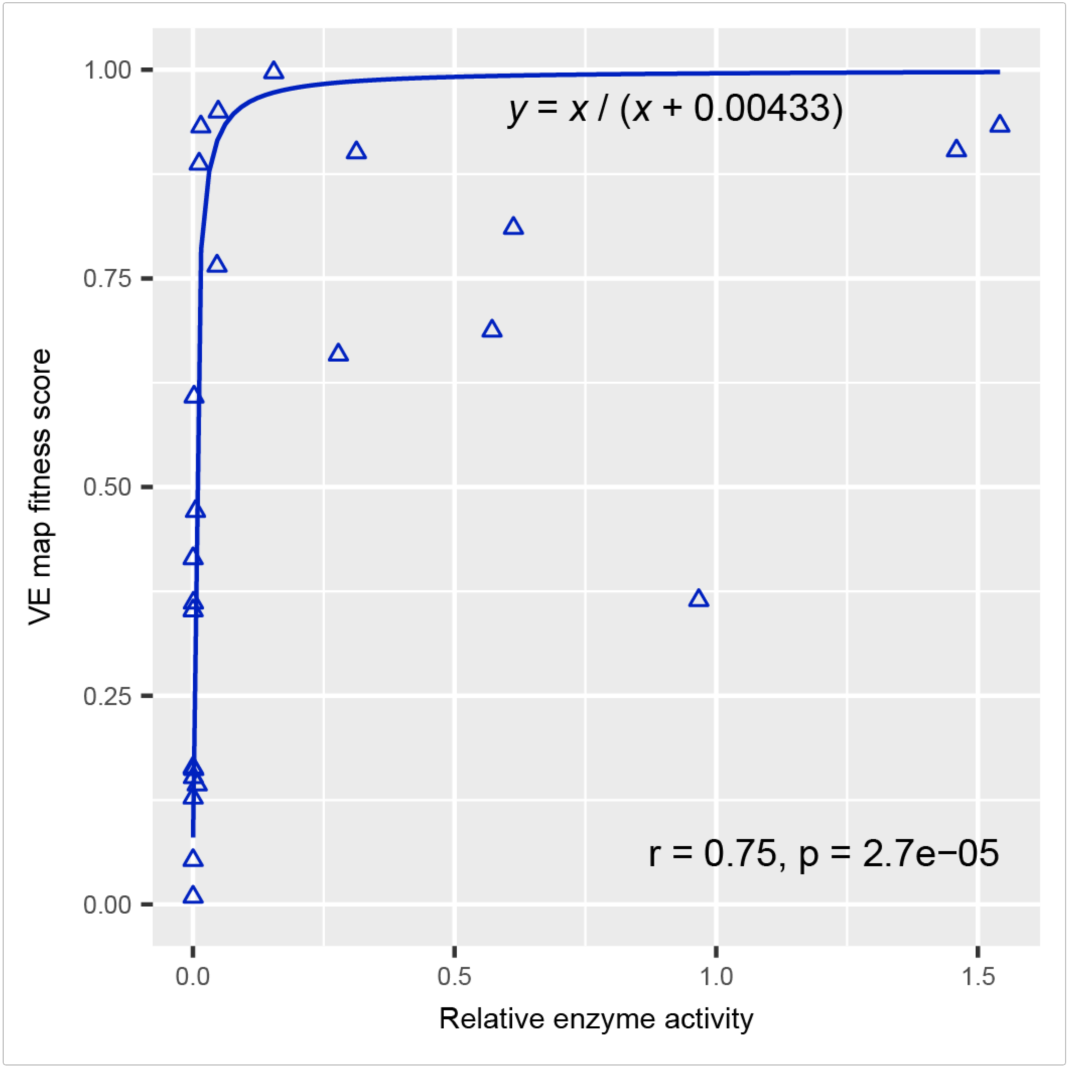
Variant effect maps show significant correlation with CBS relative enzyme activity (variant activity divided by wild type activity), following the non-linear relationship expected for recessive genes. Here, correlation (*r*) is Spearman’s rank correlation. The fitted curve was *y* = *x* / (*x* + 0.00433), where *y* is fitness score and *x* is the enzyme activity.

The well-documented clinical responsiveness to vitamin B_6_ has not yet been fully elucidated mechanistically^13,40^. To better understand the mechanistic underpinnings of vitamin B_6_ remediability of human CBS variants in the yeast model, we examined six secondary structure features, finding that positions annotated with beta-strand secondary structure tended to have lower delta scores, a trend that was modest but significant (Wilcoxon test; p = 0.0036; Supplementary Fig. 5a-f). We also found that vitamin B_6_-remediable variants tend to have higher solvent accessibility (median solvent accessibility was 82% higher in remediable variants; p = 0.00057; Supplementary Fig. 5g). Although one might naively think that variants that were the most damaging under the low vitamin B_6_ condition would be easiest to improve, the predicted change in folding energy (ΔΔG) tended to be smaller for remediable variants (median ΔΔG was 84% higher in non-remediable variants; p = 8.3 × 10^−8^; Supplementary Fig. 5h). Indeed, the strongest predictor of vitamin B_6_-remediability was having a modest fitness score in the low vitamin B_6_ map: while the median fitness score of non-remediable variants is 0, the median score of remediable variants is 0.22 (p < 2.2 × 10^−16^), indicating that some residual CBS enzyme activity is required for rescue via elevated vitamin B_6_ (Fig. 3a). This result is concordant with the clinical observation that only 9.5% of vitamin B_6_ non-responsive patients have appreciable plasma CBS activity (above 4% that of wild-type), as compared with 88% of vitamin B_6_-responsive patients having appreciable plasma CBS activity, as obtained from a study determining plasma CBS activity in homocystinuric patients by LC-MS/MS^42^ (Fig. 3b; Supplementary Table 1).

**Figure 5.**
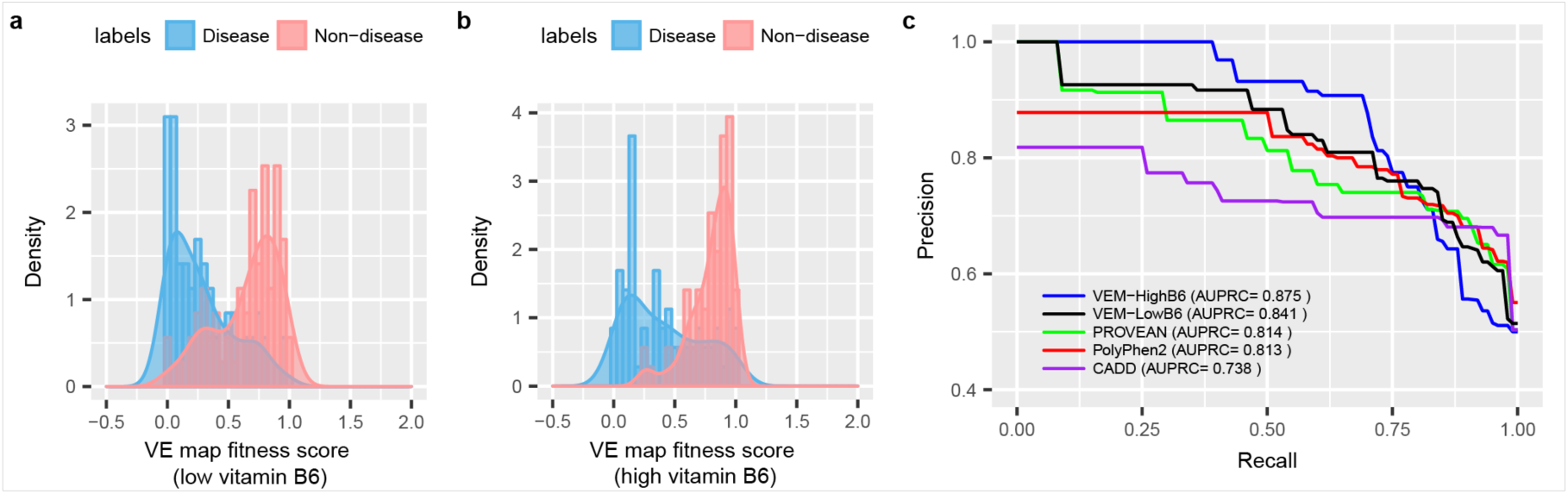
CBS variant effect maps (especially the high vitamin B_6_ map) can successfully distinguish annotated disease-causing variants from other random “non-disease” variants. (a-b) Fitness score distributions of disease and non-disease variants with low (a) or high (b) levels of vitamin B_6_. (c) Precision-recall curves for VE map fitness scores and the computational predictors PROVEAN, PolyPhen-2 and CADD capturing ability of each to discriminate disease from non-disease alleles. VE maps detect many more disease-causing variants at high precision stringency than do any of the computational methods.

To evaluate the relationship between our fitness scores and the residual CBS enzymatic activity, we examined a previous study reporting *in vitro* catalytic activities for 26 CBS missense variants expressed in *E.coli*^18^, of which 24 were well measured in our map (Supplementary Table 2). Our fitness scores exhibited a high rank correlation with measured catalytic activity (Spearman r = 0.75) with the classic nonlinear relationship between activity and fitness that was described by Kacser and Burns in 1981 to explain the molecular basis of dominant and recessive alleles^43^. Fitting a Michaelis-Menten curve relating activity to our fitness score (Materials and Methods; Fig. 4) yielded a highly-non-linear curve suggestive of the recessive behavior expected for CBS loss-of-function variants. This ‘calibration curve’ enables us to estimate the enzymatic activity of an allele given the fitness it provides in yeast.

We next assessed the potential value of our variant effect maps in identifying pathogenic CBS alleles, in terms of the trade-off between precision (fraction of predicted pathogenic variants that are annotated pathogenic) and recall (fraction of all annotated pathogenic variants that were correctly predicted). Because of the generally modest fitness scores in the C-terminal regulatory domain, and because truncation of this domain did not reduce fitness in our functional assays, we excluded CBS alleles in the regulatory domain. A set of 71 high-confidence CBS disease variants and the same number of rare variants from gnomAD were collected to evaluate prediction performance (Materials and Methods; Supplementary Table 3). Distributions of fitness scores, plotted separately for disease and non-disease alleles, clearly show that fitness scores from both low- and high-vitamin B_6_ maps can distinguish pathogenic variants (Fig. 5a-b). We then compared performance in terms of area under the precision vs recall curve (AUPRC) for our two maps with each of three computational methods (PolyPhen2, PROVEAN and CADD). Both of the variant effect maps (AUPRC=0.88 for high vitamin B_6_; AUC=0.84 for low vitamin B_6_) outperformed all three computational methods (AUC=0.81 for PolyPhen2; AUC=0.81 for PROVEAN; AUC=0.74 for CADD) (Fig. 5c). At 90% precision, the high vitamin B_6_ variant effect map captured 69% of pathogenic variants, while the best-performing computational method PROVEAN captured only 29% of pathogenic variants. These results agree with our previous study of variants in 21 human disease genes, which found that yeast complementation assays tended to have triple the sensitivity to detect pathogenic variation relative to the best computational methods^4^.

Finally, we wished to examine the ability of our maps, based on complementation phenotypes in yeast, to predict human phenotypes. For this purpose, we examined an evaluation cohort of 28 well-phenotyped homocystinuria patients (Supplementary Table 4). Two additional patients, each carrying an allele in the regulatory domain (p.Trp409*; p. Asp444Asn), were not evaluated because the yeast complementation assay does not require a functional AdoMet-binding regulatory domain (see details in Discussion). Based only on a list of alleles (blinded to all phenotypes and to the diploid genotypes of each patient), we first retrieved each allele’s low- and high-vitamin B_6_ variant effect map score and converted each fitness score to an inferred enzyme activity using the fitted curve between activity and fitness (Note: all alleles in the evaluation cohort were excluded from consideration for the Michaelis-Menten curve fitting that relates yeast fitness to enzyme activity; Supplementary Fig. 6).

**Figure 6.**
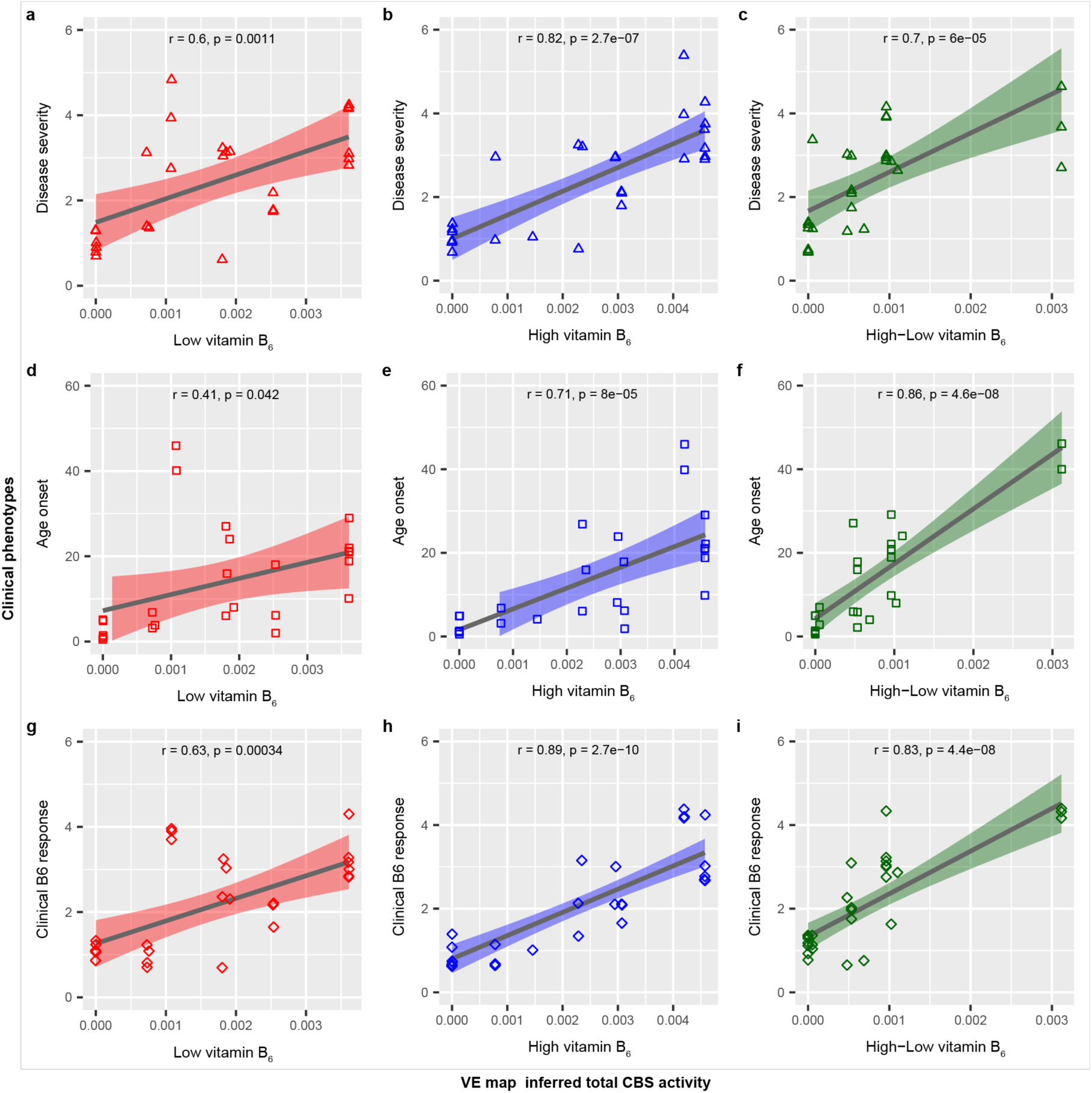
CBS VE maps, which have not been trained on patient data, successfully predict patient phenotype and response to vitamin B_6_ therapy. (a-c) Correlation between VE map-inferred total CBS activity and disease severity (d-f) Correlation between VE map-inferred total CBS activity and age onset. (g-i) Correlation between VE map-inferred total CBS activity and clinical vitamin B_6_ response. Degree of disease severity: 5-no symptoms at time of diagnosis, 4-mild disease, 3-moderate disease, 2-borderline severity, 1-severe disease. Degree of vitamin B_6_ responsiveness: 1-nonresponsive, 2-partial responsive, 3-fully responsive, 4-extremely pyridoxine responsive. A small amount of random noise (‘jitter’) was added to the categorical values of disease severity and vitamin B_6_ responsiveness to visually separate coincident data points. The amount of random noise is 0.16.

Unblinding ourselves to patient diploid genotypes, we inferred a total CBS enzyme activity for each patient by simply summing the two activity values inferred for each allele (Supplementary Table 4). Three patient CBS activity scores were calculated, corresponding to the low vitamin B_6_, high vitamin B_6_ and differential (high - low vitamin B_6_) maps. Correlation was examined for each type of patient activity score between each of three clinical phenotypes (age of onset, disease severity, and clinical response to vitamin B_6_) (Fig. 6).

Patient scores derived from the low vitamin B_6_ map yielded correlations with patient phenotypes that were significantly better than random yet still rather weak: Correlations in terms of Pearson’s r values were 0.41 (p = 0.04), 0.6 (p = 0.001) and 0.63; (p = 0.0003) for age of onset, disease severity, and clinical vitamin B_6_ response, respectively. By contrast, the high vitamin B_6_ CBS variant effect map yielded inferred patient CBS activities that correlated both strongly and significantly, with Pearson’s r = 0.71 (p = 8 × 10^−5^), 0.82 (p = 3 × 10^−7^) and 0.89 (p = 3 × 10^−10^) for age of onset, disease severity, and vitamin B_6_ responsiveness, respectively. The differential (high - low vitamin B_6_) map yielded higher correlation with age-of-onset (r = 0.86; p = 5 × 10^−8^), but reduced correlation with disease severity (r = 0.70; p = 6 × 10^−5^) and clinical vitamin B_6_ response (r = 0.83; p = 4 × 10^−8^). The performance of both high vitamin B_6_ and differential maps clearly outperformed the computational method PROVEAN, which yielded correlations of 0.44 (p = 0.03), 0.66 (p = 0.0002) and 0.65 (p = 0.0002) for age of onset, disease severity, and clinical vitamin B_6_ response, respectively.

To better understand the impact of converting our fitness scores to *in vitro* enzyme activity measurements based on the above-described calibration curve, we eliminated this step and simply summed the two yeast fitness scores corresponding to the two alleles of each patient. Eliminating the calibration step diminished our performance, but only modestly. Without this calibration, the high-vitamin B_6_ map achieved Pearson’s r = 0.70 (p = 1 × 10^−^ 4), 0.81 (p = 4 × 10^−7^) and 0.88 (p = 8.4 × 10^−10^) for age of onset, disease severity, and clinical vitamin B_6_ response, respectively (Supplementary Fig. 7). Thus, our ability to accurately predict human disease phenotypes from a CBS variant effect maps derived from yeast complementation experiments did not require calibration with *in vitro* enzyme activity assays.

In summary, variant effect maps based only on experimental measurement of growth of yeast cells expressing human *CBS* gene variants, without any computational fitting or calibration based on human traits, yielded patient activity scores that strongly correlated with clinical phenotypes in patients with classical homocystinuria.

## Discussion

Here we generated proactive maps of missense variation of the gene *CBS*. Using codon-randomizing mutagenesis to generate a mutagenized library with nearly 80% of all possible amino acid changes, we measured the functional consequences of CBS variation by measuring the effects of selection on allele frequencies during a competitive yeast complementation assay using next-generation sequencing. A machine learning model was used to impute missing data and refine the map. The resulting proactive variant effect maps agreed closely with the results of single-variant assays, and the high-vitamin B_6_ map in particular showed high performance in identifying pathogenic variants. CBS activities in individual homocystinuria patients based only on diploid *CBS* genotypes could be derived from our variant effect maps, and these correlated strongly with clinical phenotypes.

One caveat of our experimental approach is that the function of the AdoMet-binding regulatory domain was not required for functional complementation (nonsense variants truncating this domain did not reduce the ability of CBS to complement in our assay). Therefore, our assay is unsuitable for detecting some pathogenic variants that abolish AdoMet activation of the CBS enzyme activity in this regulatory domain (e.g. p.Asp444Asn) or pathogenic variants that truncate this domain (e.g. p.Trp409*). Nevertheless, other truncating variants falling within the regulatory domain did behave like null variants suggesting that our assay can still capture some large-effect variants in this domain. Given the uncertainty, however, we excluded CBS alleles in the C-terminal regulatory domain when evaluating the ability of our maps to detect pathogenic variants or infer patient phenotypes.

The clinical complications of CBS deficiency can be reduced dramatically if the diagnosis is made shortly after birth and if treatment is started in early infancy^26^. Many cases of CBS deficiency can be identified through population-level screening in newborns for the elevated methionine levels and/or methionine/phenylalanine ratio which are usually elevated in severe vitamin B_6_-non-responsive forms of homocystinurias^33,34^. However, diagnosing CBS deficiency remains a challenge, especially for the subset of vitamin B_6_-responsive patients who are most likely to benefit from vitamin B_6_ therapy (see Fig. 1a for an overview of paths for initial detection, diagnosis and therapy). Unfortunately, the majority of vitamin B_6_-responsive patients are missed on newborn screening based on detecting elevated methionine^33,34^. Although CBS deficiency can be diagnosed later in childhood upon presentation with classical symptoms of lens dislocation, skeletal abnormalities, thromboembolism and cognitive impairment, many B_6_-responsive patients do not present until adulthood^26^.

A major debate is currently underway about whether population-level genome sequencing in newborns can yield sufficient benefits, in terms of earlier detection of genetic conditions, to outweigh the costs. These costs go beyond the resources required for genome sequencing, including potential false-positive results and unsolicited findings that all lead to distress or unnecessary testing. In addition, concerns were raised about interfering with the principle of autonomy of decision when identifying variants with disease onset in the adulthood^44^. Although we can offer no overall conclusions on the advisability of population-level genome sequencing for newborns, it is worth considering the potential value that population-level screening by genome sequencing might bring to each of many individually-rare genetic diseases.

We found that our *CBS* variant effect map accurately distinguished annotated pathogenic variants from unannotated variants. At a stringent threshold achieving 90% precision in our test set, the variant effect map captured more than twice the number of pathogenic variants than did the best-performing computational prediction method at the same 90% precision stringency. The map enabled predictions of CBS activity in human patients, which we assessed using a cohort of 28 subjects. Although larger cohorts would provide more precise performance estimates, map-derived predictions of patient activity correlated strikingly well with patient phenotype and therapeutic response. This success was achieved by simply summing map-predicted activities for the two patient alleles. Future performance might be improved by testing allele combinations in a compound heterozygous diploid model system. Already, however, the predictive power suggests potential clinical uses of the *CBS* variant effect map. For example, genome analysis of patients with cognitive defects, for which many causes are possible *a priori*, may reveal unexpected *CBS* variants that would currently be classified as VUS. In this scenario, genomic sequencing coupled with the VE map could sensitively detect deleterious CBS variants, and thus trigger tHcy measurement and further confirmatory testing, while reducing false positives. Should population-level newborn genome sequences become available in the future, genome interpretation using the *CBS* variant effect map may trigger testing for homocystinuria in still-asymptomatic patients. Such testing, occurring even in the absence of elevated methionine or early childhood symptoms, could enable earlier identification of many additional cases of CBS deficiency, particularly those that would be most responsive to vitamin B_6_-therapy.

There are 497 human genes that encode a cofactor-dependent enzyme, of which at least 193 (39%) reportedly harbor disease-causing variants (http://au.expasy.org/enzyme; Supplementary Table 5). Based on overall rates of missense variation^45,46^, we might expect every individual to carry roughly 5-10 missense alleles in these enzymes on average. According to a recent survey of assayable genes^9^, 53% of these genes have assays tractable for VE-mapping and ∼10% have a yeast complementation assay. Thus, our study provides a blueprint for systematically and proactively cataloguing missense variant effects and their dependence on co-factor levels for this important class of human enzymes.

Despite the ability of cell-based complementation assays to detect deleterious variants with high accuracy, only with additional context can they explain the mechanism of the defect. For example, it is unclear whether protein function has been reduced due to a direct reduction in enzymatic activity, disruption of the ability to receive an activating modification, or due to misfolding that reduces stability and leads to a lower steady-state protein expression level. There is now ample precedent for VE maps that measure the effect of variation on ‘sub-functions’ such as protein interaction (which might include tetramerization for CBS), protein abundance^47^ or post-translational modification.

## Materials and Methods

### Strains and plasmids

The *S. cerevisiae* strain (*MATα cys4Δ∷KanMX his3Δ1 leu2Δ0 lys2Δ0 ura3Δ0*), used as a host for the CBS variant library, was derived from the yeast knockout collection^48^. The Gateway destination vector pAG415GAL-ccdB (CEN/ARS-based, *GAL1* promoter, and *LEU2* marker) was purchased from Addgene and served as the yeast expression vector. The CBS ORF clone was from Human ORFeome v8.1 library^49^.

### Construction of *CBS* variant library by POPcode mutagenesis

A library of CBS variants was constructed using an oligo-pool based saturation mutagenesis method (<u>P</u>recision <u>O</u>ligo-<u>P</u>ool based <u>Code</u> Alteration or POPCode). Details are described below with some technical advancements that decrease the frame-shift mutation rate and therefore render the method suitable for mutagenizing larger genes. Oligonucleotides of 28-38 bases were designed to target each codon in the open reading frame of *CBS*, such that the targeted codon is replaced with a NNK_-_degenerate codon (a mixture of all four nucleotides in the 1^st^ and 2^nd^ codon positions, and a mixture of G and T in the 3^rd^ position). The 550 oligos were synthesized and combined into a single pool. The uracilated wild type template was generated by PCR-amplifying the ORF with dNTP/dUTP mix and KAPA HiFi Uracil+ DNA polymerase (KapaBiosystems) and mixed with the phosphorylated oligonucleotide pool. Oligos were annealed to the template by heating the mixture to 95 degrees C for 3 minutes and then cooled to 4 degrees C. Gaps between annealed oligonucleotides were then filled with KAPA HiFi Uracil+ DNA polymerase followed by nick-sealing with T4 DNA ligase (*NEB*). After degradation of the uracil-doped wild-type strand using Uracil-DNA-Glycosylase (UDG) (*NEB*), the mutated strand was amplified with attB-sites-containing primers and subsequently transferred *en masse* to a Donor vector via the Gateway BP reaction to generate a library of Entry clones. To enable yeast expression, the library was further transferred to pAG415-ccdB by *en masse* Gateway LR reaction and transformed into the *S. cerevisiae cys4* mutant strain. To maintain the complexity of the library, plasmids were purified from >100,000 clones at each transferring step and ∼1,000,000 yeast transformants were pooled to form the host library.

### High-throughput yeast-based complementation screen

The yeast based functional complementation assay for CBS function has been well established for characterizing individual variants^35,36,39^. Details are provided here for high-throughput complementation screening: Plasmids extracted from a pool of >100,000 *E. coli* clones were transformed into the *S. cerevisiae cys4* mutant strain yielding ∼1M total transformants. Plasmids were prepared from two of ∼1×10^8^ of cells and used as templates for the downstream tiling PCR (two replicates of non-selective condition). Selective media was made with yeast nitrogen base lacking all vitamins and amino acids (USBiological). All other vitamins except vitamin B_6_ were added back at standard concentrations and vitamin B_6_ was supplemented at three different concentrations: 0, 1, and 400 ng/ml. Histidine, uracil and lysine were added to relieve auxotrophies in the mutant strain and 2% galactose was used as carbon source to induce *GAL1* promoter-driven expression. For each of the three pooled complementation assays (each using a different concentration of vitamin B_6_), ∼4×10^8^ of cells were inoculated into 200ml selective medium for each of two replicates. In parallel, plasmid expressing the wild-type ORF was similarly transformed to the *S. cerevisiae cys4* mutant strain in selective media. Each culture (with two biological replicate cultures for both the selective and non-selective condition) was grown to full density (5-6 doublings) with shaking at 30°C. Plasmids extracted from ∼1×10^8^ of cells of each culture were used as templates for the downstream tiling PCR.

### TileSeq for mapping functional variation

For each plasmid library, the tiling PCR was performed in two steps: (i) the targeted region of the ORF was amplified with primers carrying a binding site for Illumina sequencing adaptors, (ii) each first-step amplicon was indexed with an Illumina sequencing adaptor in the second-step PCR. We perform paired-end sequencing on the tiled regions across the ORF with an average sequencing depth of ∼2 million reads. All raw sequencing reads were mapped to CBS using bowtie2^50^ to generate alignment files for both the forward and reverse reads. A custom Perl script (https://bitbucket.org/rothlabto/tileseq_package) was used to parse the alignment files to count the number of co-occurrences of a codon change in both paired reads and the mutational counts for each tiled region were subsequently normalized by the corresponding sequencing depth. Then, the normalized mutational counts from the wild type control libraries (which enable adjustment for inflated allele frequencies due to sequencing errors) were subtracted from the normalized mutational counts from the non-selective and selective conditions respectively. Finally, an enrichment ratio (Φ) was calculated for each mutation based on the adjusted mutation counts before and after selection.

### Calculation of fitness scores and remediation scores

A fitness score (s_MUT_) was calculated for each variant as ln(Φ_MUT_/Φ_STOP_)/ln(Φ_SYN_/Φ_STOP_), where Φ_MUT_ is the enrichment ratio calculated for each variant, Φ_STOP_ is the median enrichment ratio of all nonsense variants and Φ_SYN_ is the median enrichment ratio of all synonymous variants, such that s_MUT_ equals zero when Φ_MUT_ equals Φ_STOP_ and s_MUT_ equals one when Φ_MUT_ equals Φ_SYN_. Well-measured variants were selected by applying two filters: allele frequency is greater than 0.005% and standard error is less than 0.2. A remediation score was calculated as the difference between fitness scores at high (400 ng/ml) and low (both 0 and 1 ng/ml, with fitness scores averaged due to high agreement between these screens) vitamin B_6_ concentrations.

A MAP (maximum *a posteriori*) estimate of the error (*σ*) in each fitness score was derived via a weighted average of the observed variance and the *a priori* estimate of *σ*. We used 6 pseudocounts, so that the observed variance was given weight 2/(2+6), based on having two replicates, and the prior variance was given weight 6/(2+6). The prior estimate of *σ* is based on an overall regression of *σ* values against fitness values^51^.

To produce a complete variant effect map, missing values were estimated by imputation (unpublished result). Briefly, the imputation machine learning model was trained on the fitness scores of the experimentally well-covered variants using the Gradient Boosted Tree (GBT) method. The features used in the model included precomputed PolyPhen-2 and PROVEAN scores, chemical and physical properties of the wild type and substituted amino acids, protein structure-related information, and the average fitness value at each position. Final variant effect maps use scores that were refined using the weighted average of imputed and measured values (weighting by the inverse-square of estimated standard error in each input value). Although we used the original well-measured values for all variant effect map performance analysis, the results were similar using the imputed and refined map values.

To estimate agreement with previous individual yeast complementation assay data^39^, only well-measured values were used (only 10 out of 78 variants were poorly measured and therefore excluded for this analysis). All other analyses used the final imputed and refined map, although we note that all patient missense variants happened to be well-measured so that some values were refined but none were imputed.

### Classification of vitamin B_6_ remediable and non-remediable deleterious variants

Using the fitness score distribution of all synonymous variants as an empirical null distribution, FDR-adjusted p-values were assigned to all missense variants. The fitness score corresponding to FDR = 5% was determined to be 0.45, so that missense variants for which the upper end of the 95% confidence interval of their fitness scores was less than 0.45 were classified as deleterious variants. Then, for each deleterious variant, a delta fitness score (high vitamin B_6_ – low vitamin B_6_) was calculated. Using the delta fitness score distribution of all nonsense variants as an empirical null distribution, FDR-adjusted p-values were assigned to all missense variants and a delta fitness score threshold (0.29, corresponding to FDR = 5%) was used to identify B_6_-remediable variants. Missense variants for which the lower end of the 95% confidence interval of their delta fitness scores was greater than 0.29 were classified as B_6_ remediable.

### Investigation of relationship between fitness score and enzyme activity

Biochemical properties including enzyme activity have been examined for 27 CBS variants expressed in *E.coli* in a previous study^18^, among which 24 missense variants that were well measured in this study were selected to investigate the relationship between the fitness score and the experimentally determined enzyme activity. The best correlation was observed between fitness score with high level of vitamin B_6_ and the relative CBS enzyme activity (variant activity divided by wild type activity) with AdoMet at 37°C. A Michaelis-Menten curve (of the form *y* = *x* / (*x*+*k*), where *y* is fitness score, *x* is relative enzyme activity, and *k* is a constant) was fitted to describe the non-linear relationship between fitness and activity.

### Selection of disease and non-disease associated variants

To define a set of disease-associated CBS variants, we considered 86 unique missense variants in the CBS mutation database^17^ that were not linked to a secondary mutation, then reviewed the relevant literature, accepting only the 79 disease variants that we considered to be high confidence (after excluding variants in the regulatory domain, 71 disease variants was selected for prediction performance evaluation; Supplementary Table 3). For non-disease-associated variants, we selected all CBS missense variants deposited in gnomAD^1^ that meet the following criteria: (i) there is no disease association or experimental evidence of functional impact and (ii) the variant was observed at least twice (Supplementary Table 3). All CBS variants from gnomAD that met these criteria were rare, with minor allele frequency less than 0.005.

### Patient phenotype data collection

All patients have been followed in the Metabolic Center in the Department of Pediatrics and Adolescent Medicine, the General University Hospital in Prague. The clinical, biochemical and molecular genetic data were obtained during routine care, patients gave their informed consent for DNA analysis. Plasma CBS activity was measured within a research project after obtaining patient informed consent, which included also a consent with publication of clinical, enzymatic and molecular genetic data (approval of the Ethics Committee 1194/13 S-IV).

To assess the clinical severity and vitamin B_6_-responsiveness of CBS deficiency, we developed a semi-quantitative scoring system based both on tHcy changes after vitamin B_6_ administration and on the need for additional therapy. Nonresponsive patients, needing low methionine diet and betaine (regardless of vitamin B_6_ therapy), were assigned vitamin B_6_-responsiveness score 1. Partially responsive patients, needing both large doses of vitamin B_6_ and low methionine diet, were given score 2. Fully responsive patients needing only vitamin B_6_ in a dose above 0.5 mg/kg/day to yield tHcy < 50 µmol/L received score 3. Extremely responsive patients, needing B_6_ at a dose below 0.5 mg/kg/day to yield tHcy < 50 µmol/L, were given a vitamin B_6_-responsiveness score of 4.

Disease severity was scored according to the presence of typical clinical complications at the time of diagnosis or during follow-up in poorly compliant patients, and could not have been determined in two patients detected by newborn screening. Patients showing no symptoms at the time of diagnosis (i.e. detected by screening family members of patients with diagnosed CBS deficiency) received severity score 5; Patients with mild disease (thrombosis in any vascular bed with no other symptoms) received score 4. Patients with moderate disease (connective tissue involvement with or without thrombosis) were assigned score 3. Those with borderline severity (mild cognitive impairment with good social outcome, regardless of other somatic complications) were given score 2. Severe disease patients (having severe neuropsychiatric complications including poor social outcome, regardless of other somatic complications) were defined to have severity score 1.

## Supporting information

## Acknowledgements

The authors thank Drs. Andreas Schulze and Henricus Blom for helpful discussion, Dr. Jakub Krijt and Ms. Jitka Sokolová, MSc. for analyzing plasma CBS activity and Ms. Hana Vlášková for molecular genetic testing. This work was supported in part by the One Brave Idea Initiative, the National Human Genome Research Institute of the National Institutes of Health (NIH/NHGRI) Center of Excellence in Genomic Science (CEGS) Initiative (HG004233), the Canadian Excellence Research Chairs (CERC) Program, a Canadian Institutes of Health Research Foundation Grant, the Canada Foundation for Innovation, by RVO-VFN 64165 (General University Hospital in Prague) and Progres Q26 (Charles University).

