## Supplementary material for "A proactive genotype-to-patient-phenotype map for cystathionine beta-synthase"

### Supplementary Figures

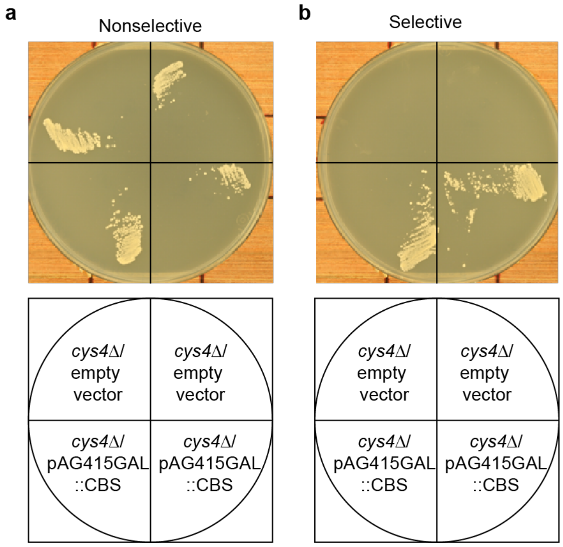


Supplementary Figure 1. Expression of human CBS protein can rescue growth of yeast cys4Δ strain in the absence of exogenous cysteine. Growth of cys4Δ strain expressing CBS gene or carrying empty vector in non-selective (a) or selective (b) medium. The nonselective medium is synthetic complete medium lacking cysteine and supplemented with glutathione, a stable source of cysteine. The selective medium is synthetic complete medium lacking cysteine and glutathione. All medium was supplemented with galactose as carbon source to induce the expression.

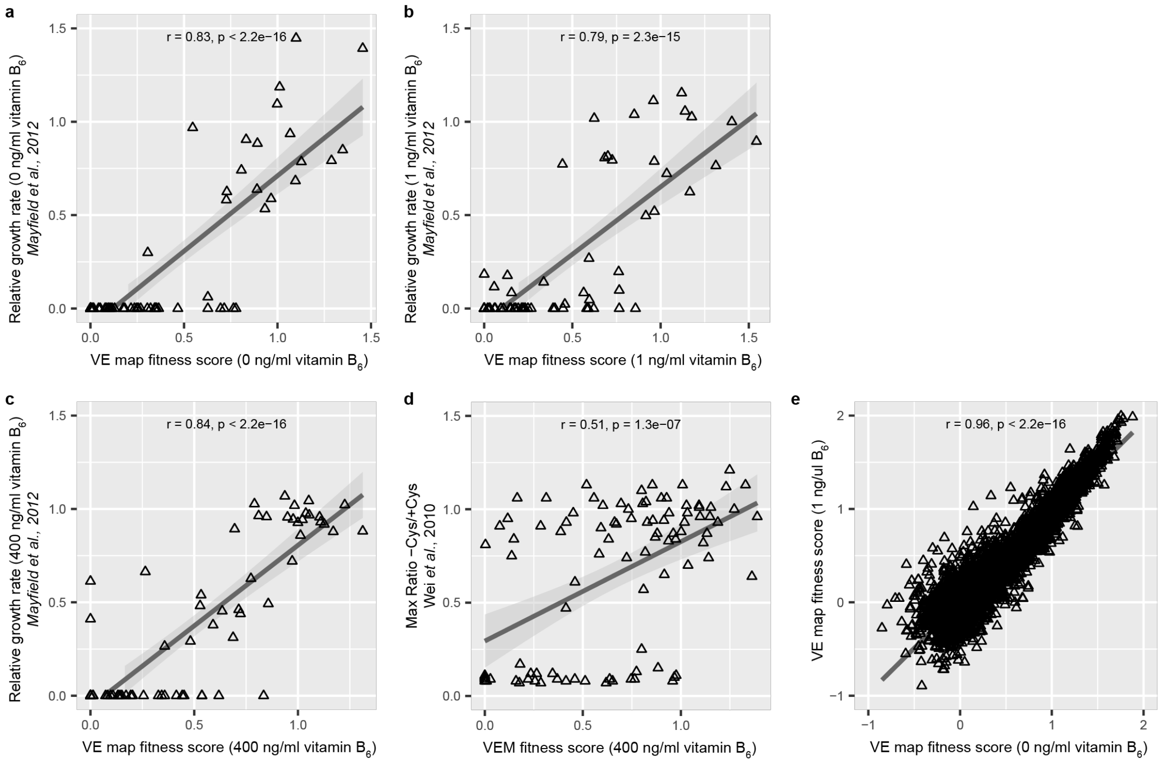


Supplementary Figure 2. VE map fitness scores are in good agreement with relative growth rates determined in single-variant assays and strong correlation between VE map fitness scores with low concentrations of vitamin $B_6$. (a-c) Correlation between fitness scores from VE maps and relative growth rates from single-variant assays at three different vitamin $B_6$ concentrations. (d) Correlation between fitness scores from high vitamin B6 VE map and Cys-/Cys+ growth ratios from single-variant assays. (e) Correlation between VE map fitness scores with 0 and 1 ng/ml vitamin $B_6$. The correlation test is Pearson correlation.

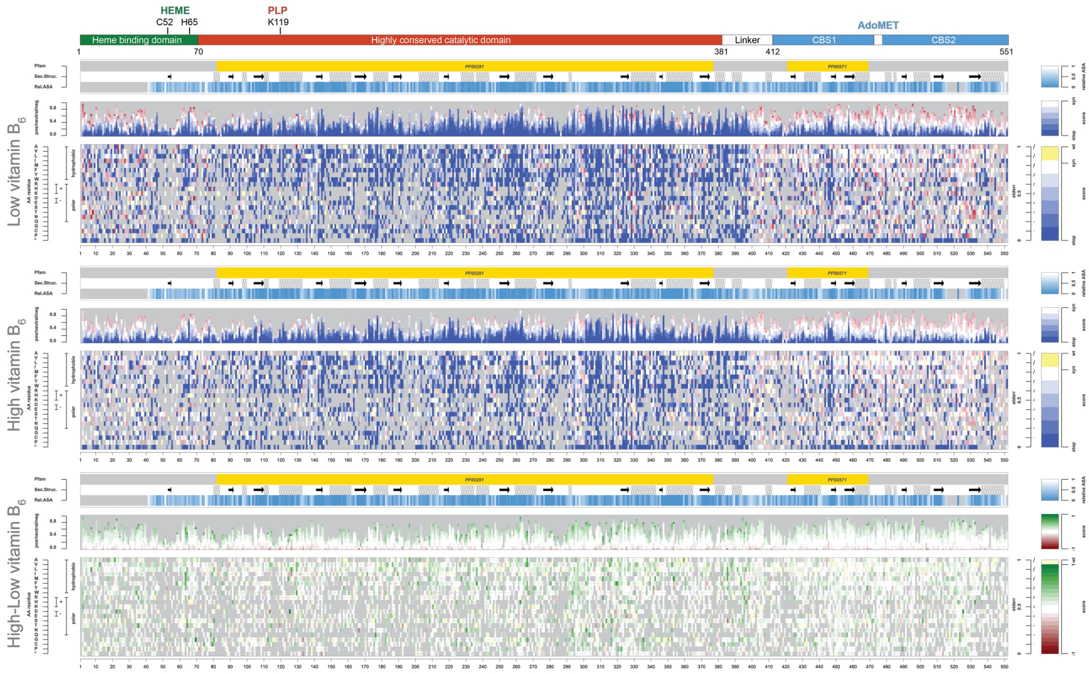


Supplementary Figure 3. VE maps for CBS before computational imputation and refinement: fitness landscape with low level of vitamin $B_6$ (top), fitness landscape with high level of vitamin $B_6$ (middle) and delta fitness (high-low vitamin $B_6$) landscape (bottom). For the two fitness landscapes (high or low vitamin $B_6$), a functional score of 0 (blue) corresponds to a fitness equivalent to the median fitness of stop codon variants. A score of 1 (white) corresponds to a fitness equivalent to the median fitness of synonymous variants. A score greater than 1 (red) corresponds to fitness above the median fitness of synonymous variants. For the delta fitness landscape (high-low vitamin$B_6$), substitutions were colored green if delta fitness score is positive and red if negative.

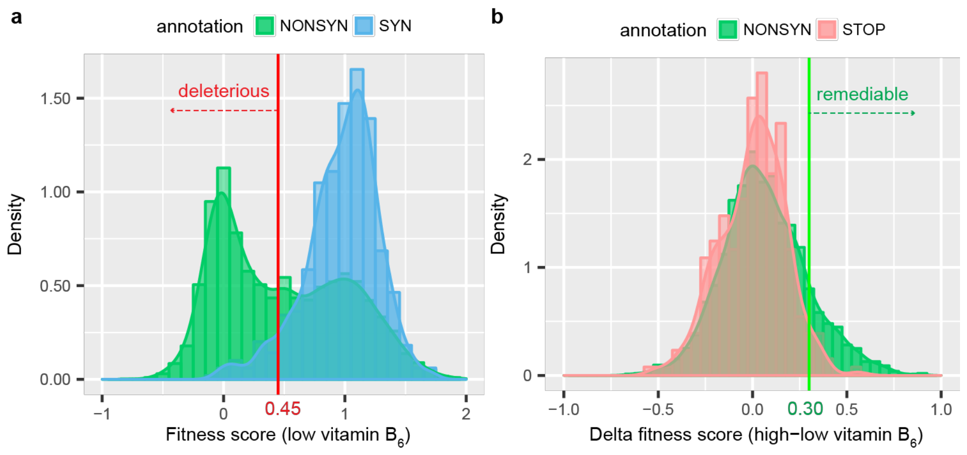


Supplementary Figure 4. Determination of fitness score and delta fitness score cutoff values for significant fitness defect (0.45) and vitamin $B_6$ remediability (0.30). (a) Classification of deleterious CBS variants with significant fitness defect using fitness score (low vitamin $B_6$) distribution of synonymous variants as the null distribution. (b) Classification of vitamin $B_6$-remediable deleterious variants using delta fitness (high-low vitamin $B_6$) distribution of stop codon variants as the null distribution.

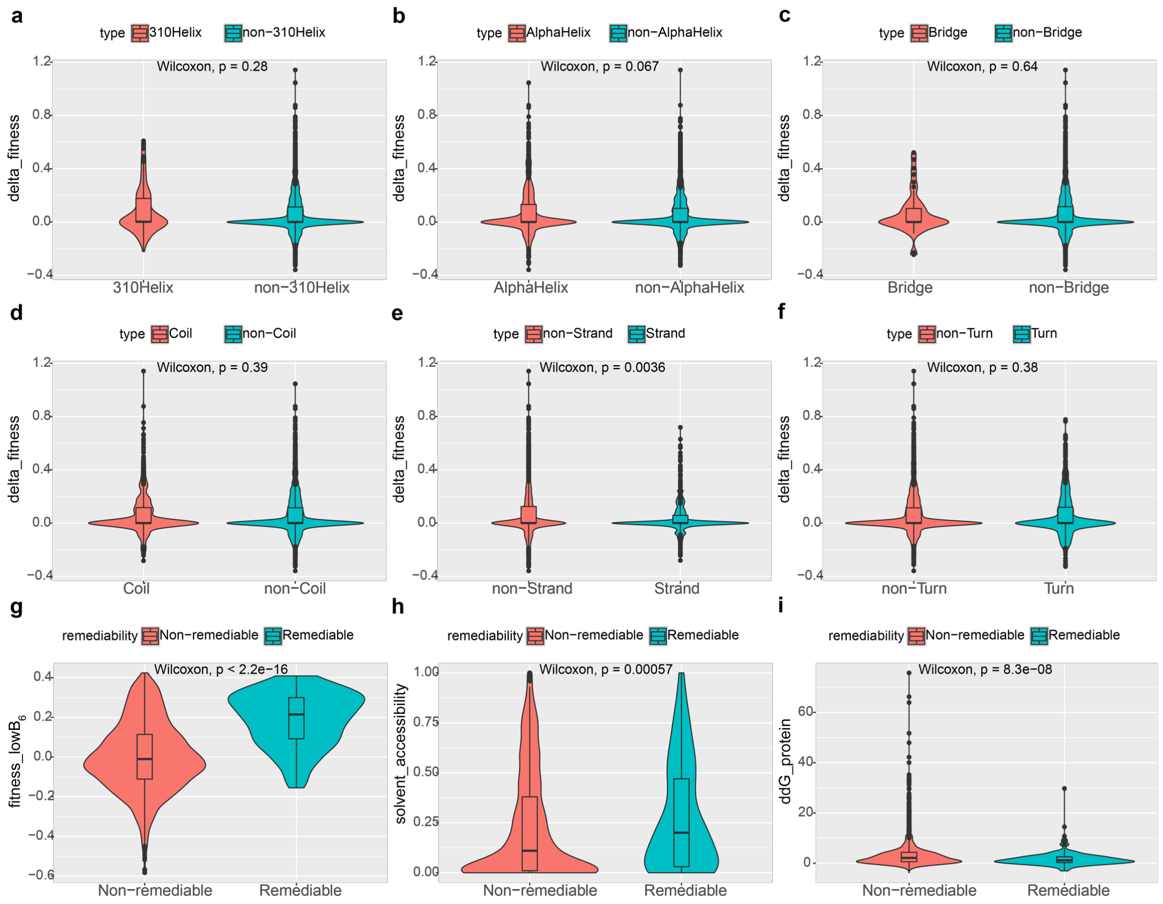


Supplementary Figure 5. Vitamin $B_6$-remediable variants tend to have non-beta-strand secondary structures, higher solvent accessibility, smaller change in folding energy and higher fitness score in low vitamin $B_6$ map. Feature analysis for vitamin $B_6$ remediability of CBS variants. (a-f) Delta fitness score distribution comparison between different secondary structural features: 310Helix/non-310Helix (a), AlphaHelix/non-AlphaHelix (b), Bridge/non-Bridge (c), Coil/non-Coil (d), β-strand/non-β-Strand (e), Turn/non-Turn (f). (g-i) Comparison between vitamin $B_6$-remediable and vitamin $B_6$-non-remediable variants in terms of their (g) solvent accessibility, free energy change (h) and fitness score (i) distributions.

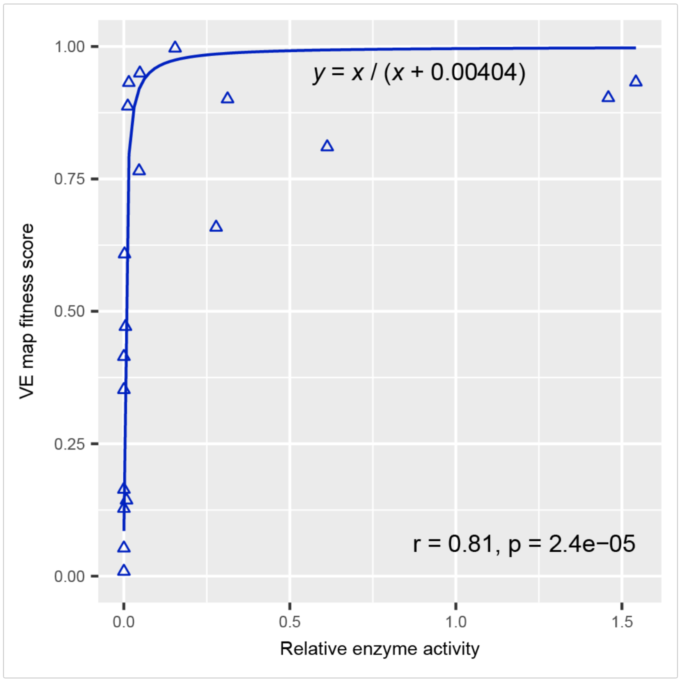


Supplementary Figure 6. Variant effect maps show significant correlation with CBS relative enzyme activity (variant activity divided by wild type activity), following the non-linear relationship expected for recessive genes (CBS alleles observed in the homocystinuria cohort were excluded). The correlation test is Spearman's rank correlation. The fitted curve is y = x/(x+0.00404), where y is fitness score, x is the enzyme activity. This curve was used to convert VE map fitness score to enzyme activity to predict patients' clinical phenotypes in the homocystinuria cohort.

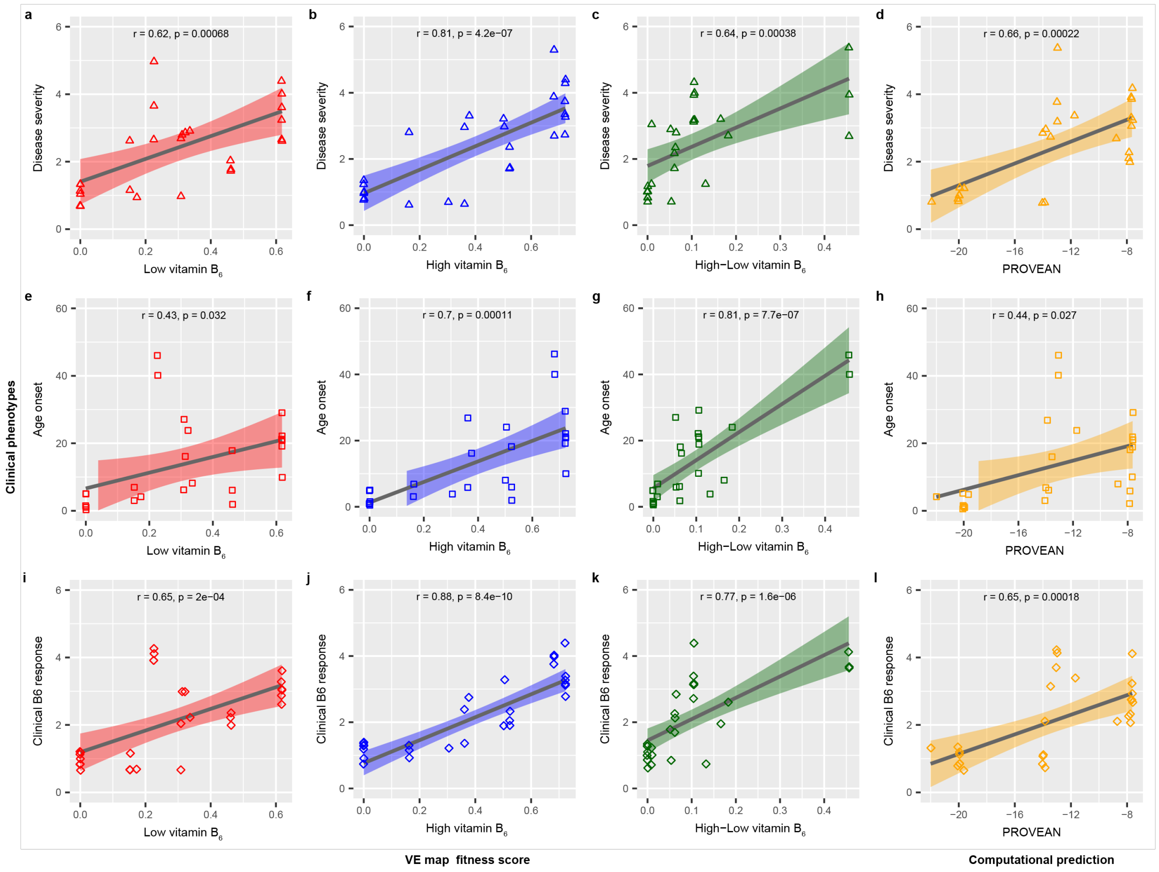


Supplementary Figure 7. CBS VE map fitness scores (not converted to enzyme activity) successfully predict patient phenotype and response to vitamin $B_6$ therapy, and outperform computational prediction. (a-c) Correlation between VE map inferred fitness scores and disease severity. (e-g) Correlation between VE map inferred fitness scores and age onset. (i-k) Correlation between VE map inferred fitness scores and clinical $B_6$ response. (d,h,l) Correlation between PROVEAN scores and three clinical phenotypes. The correlation test is Pearson correlation. Degrees of disease severity: 5-no symptoms at time of diagnosis, 4-mild disease, 3-moderate disease, 2-borderline severity, 1- severe disease. Degrees of vitamin $B_6$ responsiveness: 1-nonresponsive, 2-partial responsive, 3-fully responsive, 4-extremely pyridoxine responsive. A small amount of random noise (‘jitter’) was added to the categorical values of disease severity and vitamin $B_6$ responsiveness to visually separate coincident data points. The amount of random noise is 0.16.

### Supplementary Tables

Supplementary Table 1. Plasma CBS activity in vitamin B6 responder and non-responder.

| patient_ID | vitamin $B_6$ response | relative activity to median of controls |
|---|---|---|
| CZ-1 | $B_6$-nonresponder | 0.0% |
| CZ-2 | $B_6$-nonresponder | 0.0% |
| CZ-3 | $B_6$-responder | 4.0% |
| CZ-4 | $B_6$-responder | 15.0% |
| CZ-5 | $B_6$-responder | 0.0% |
| CZ-6 | $B_6$-responder | 4.0% |
| CZ-7 | $B_6$-responder | 16.0% |
| CZ-8 | $B_6$-responder | 22.0% |
| CZ-9 | $B_6$-responder | 4.0% |
| CZ-10 | $B_6$-responder | 6.0% |
| E-1 | $B_6$-nonresponder | 2.0% |
| E-2 | $B_6$-nonresponder | 4.0% |
| E-3 | $B_6$-nonresponder | 2.0% |
| E-4 | $B_6$-nonresponder | 3.0% |
| E-5 | $B_6$-nonresponder | 0.0% |
| E-6 | $B_6$-nonresponder | 1.0% |
| E-7 | $B_6$-nonresponder | 2.0% |
| E-8 | $B_6$-nonresponder | 1.0% |
| E-9 | $B_6$-nonresponder | 1.0% |
| E-10 | $B_6$-nonresponder | 2.0% |
| E-11 | $B_6$-nonresponder | 1.0% |
| E-12 | $B_6$-nonresponder | 1.0% |
| E-13 | $B_6$-nonresponder | 4.0% |
| E-16 | $B_6$-nonresponder | 1.0% |
| E-17 | $B_6$-nonresponder | 1.0% |
| E-18 | $B_6$-nonresponder | 3.0% |
| E-19 | $B_6$-nonresponder | 1.0% |
| E-20 | $B_6$-nonresponder | 0.0% |
| E-21 | $B_6$-nonresponder | 2.0% |

Data was collected from Alcaide *et al* 2015, reference 42.

Supplementary Table 2. Relative In-vitro catalytic activity for 24 CBS missense variants expressed in *E. coli* and their corresponding high vitamin $B_6$ VE map fitness scores.

| Mutant | VE map fitness (high vitamin $B_6$) | Relative CBS enzyme activity* | | |
|---|---|---|---|---|
| | | No addition at 37°C | AdoMet at 37°C | AdoHcy at 37°C |
| C165Y | 0.1521 | 0.0077 | 0.0011 | 0.0047 |
| A114V | 0.6873 | 0.7687 | 0.5717 | 0.6113 |
| R369C | 0.8873 | 0.0175 | 0.0121 | 0.0125 |
| T191M | 0.0529 | 0.0028 | 0.0004 | 0.0010 |
| S466L | 0.8104 | 2.4261 | 0.6126 | 1.8480 |
| G148R | 0.4145 | 0.0014 | 0.0004 | 0.0016 |
| P422L | 0.9009 | 0.4590 | 0.3125 | 0.3501 |
| P49L | 0.3644 | 1.0371 | 0.9666 | 0.8929 |
| T262R | 0.009 | 0.0014 | 0.0004 | 0.0010 |
| N228K | 0.3521 | 0.0014 | 0.0004 | 0.0010 |
| E144K | 0.1619 | 0.0042 | 0.0017 | 0.0026 |
| K102N | 0.9497 | 0.0617 | 0.0484 | 0.0481 |
| R266K | 0.9969 | 0.1773 | 0.1546 | 0.1536 |
| L539S | 0.6082 | 0.0084 | 0.0023 | 0.0084 |
| G307S | 0.1277 | 0.0021 | 0.0004 | 0.0021 |
| R439Q | 0.9031 | 1.1724 | 1.4590 | 0.9394 |
| E176K | 0.4711 | 0.0343 | 0.0051 | 0.0230 |
| H65R | 0.1434 | 0.0392 | 0.0083 | 0.0099 |
| G305R | 0.1637 | 0.0014 | 0.0004 | 0.0010 |
| E302K | 0.6586 | 0.9544 | 0.2780 | 0.5867 |
| D444N | 0.9327 | 1.6377 | 1.5419 | 1.6385 |
| I278T | 0.3613 | 0.0028 | 0.0011 | 0.0021 |
| R125Q | 0.9318 | 0.0161 | 0.0153 | 0.0141 |
| V180A | 0.7651 | 0.0918 | 0.0462 | 0.1134 |

*The relative enzyme activity is calculated as variant activity divided by wild type activity. Data on mutant enzymes expressed in *E. coli* taken from Kožich et al. 2010, reference 18.

Supplementary Table 3. Selected CBS variants and prediction scores for evaluation of prediction performance.

| AAchange | wt | pos | mut | VE map fitness (low vitamin B6) | VE map fitness (high vitamin B6) | PROVEAN | PPH2 | CADD | MAF | labels |
|---|---|---|---|---|---|---|---|---|---|---|
| P2L | P | 2 | L | 0.7598 | 0.8985 | -0.62 | 0.011 | 13.98 | 4.46E-05 | neutral |
| P6S | P | 6 | S | 0.6471 | 0.7618 | 0.12 | 0.002 | 0.008 | 1.48E-05 | neutral |
| G11R | G | 11 | R | 0.8445 | 0.9934 | -0.04 | 0.005 | 10.24 | 1.25E-05 | neutral |
| P12S | P | 12 | S | 0.7752 | 0.9054 | 0.11 | 0 | 0.001 | 8.31E-06 | neutral |
| C15F | C | 15 | F | 0.8968 | 0.895 | -1.95 | 0.846 | 24.7 | 2.90E-05 | neutral |
| H17L | H | 17 | L | 0.929 | 0.9419 | 0.04 | 0.002 | 12.25 | 2.90E-05 | neutral |
| S19L | S | 19 | L | 0.369 | 0.4252 | -0.46 | 0.002 | 10.35 | 1.10E-05 | neutral |
| H22R | H | 22 | R | 0.8072 | 0.9834 | -0.38 | 0.001 | 0.002 | 3.66E-05 | neutral |
| S23L | S | 23 | L | 0.7938 | 0.818 | -0.2 | 0 | 0.412 | 1.24E-05 | neutral |
| A24V | A | 24 | V | 0.8504 | 0.8744 | -0.53 | 0.002 | 0.08 | 3.71E-05 | neutral |
| S27R | S | 27 | R | 0.2515 | 0.6158 | -0.51 | 0.001 | 6.248 | 8.21E-06 | neutral |
| S27I | S | 27 | I | 0.5791 | 0.7905 | -0.56 | 0.086 | 12.1 | 8.21E-06 | neutral |
| S27N | S | 27 | N | 0.7224 | 0.8328 | -0.21 | 0 | 4.361 | 1.23E-05 | neutral |
| S32P | S | 32 | P | 0.8583 | 0.9421 | -0.34 | 0 | 5.441 | 1.09E-05 | neutral |
| P41S | P | 41 | S | 0.7628 | 0.9964 | 0.87 | 0 | 3.693 | 8.14E-06 | neutral |
| R45Q | R | 45 | Q | 0.5558 | 0.931 | -2.32 | 0.625 | 23.2 | 8.55E-05 | neutral |
| R45W | R | 45 | W | 0.2801 | 0.7134 | -4.95 | 0.998 | 29.1 | 5.96E-04 | neutral |
| P49L | P | 49 | L | 0.1763 | 0.3644 | -8.11 | 0.977 | 25.7 | 1.48E-04 | deleterious |
| R51S | R | 51 | S | 0.9379 | 0.8914 | -3.24 | 0.223 | 24.2 | 6.46E-05 | neutral |
| P59S | P | 59 | S | 0.9924 | 0.8968 | 0.06 | 0 | 0.151 | 4.49E-05 | neutral |
| E62K | E | 62 | K | 0.8049 | 0.8577 | -0.54 | 0.007 | 8.276 | 1.81E-05 | neutral |
| H65R | H | 65 | R | 0.409 | 0.1434 | -6.8 | 0.99 | 23.4 | 4.08E-06 | deleterious |
| P70L | P | 70 | L | 0.5804 | 0.6709 | -0.22 | 0 | 0.004 | 4.35E-05 | neutral |
| S73C | S | 73 | C | 0.7138 | 0.7706 | -2.87 | 0.289 | 15.43 | 2.04E-05 | neutral |
| P74L | P | 74 | L | 0.949 | 0.9605 | -4.4 | 0.062 | 21.3 | 1.75E-04 | neutral |
| K75N | K | 75 | N | 0.2381 | 0.7487 | -2.9 | 0.025 | 19.19 | 8.54E-05 | neutral |
| G85R | G | 85 | R | 0.74 | 0.9801 | -7.76 | 1 | 32 | 4.06E-06 | deleterious |
| T87N | T | 87 | N | 0.4798 | 0.5161 | -4.71 | 1 | 26.5 | NA | deleterious |
| P88S | P | 88 | S | 0.3401 | 0.6333 | -7.76 | 1 | 26.8 | NA | deleterious |
| G100S | G | 100 | S | 0.9729 | 0.9437 | -5.13 | 0.974 | 32 | 1.08E-05 | neutral |
| L101V | L | 101 | V | 0.4019 | 0.541 | -1.89 | 0.488 | 16.52 | 1.22E-05 | neutral |
| L101P | L | 101 | P | 0.4989 | 0.4233 | -6.2 | 1 | 27.4 | NA | deleterious |
| C109R | C | 109 | R | 0.646 | 0.4458 | -11.9 | 1 | 27.5 | 1.63E-05 | deleterious |
| A114S | A | 114 | S | 0.7893 | 0.8138 | -2.05 | 0.206 | 28.7 | 1.22E-05 | neutral |
| A114T | A | 114 | T | 0.3535 | 0.8059 | -2.82 | 0.992 | 35 | 1.22E-05 | neutral |
| A114V | A | 114 | V | 0.3299 | 0.6873 | -3.15 | 0.969 | 33 | 0.0002132 | deleterious |
| G116R | G | 116 | R | 0.2566 | 0.419 | -7.95 | 1 | 35 | NA | deleterious |

| | | | | | | | | | | |
|---|---|---|---|---|---|---|---|---|---|---|
| V118M | V | 118 | M | 0.8191 | 0.8578 | -2.76 | 0.999 | 34 | 1.81E-05 | neutral |
| R121C | R | 121 | C | 0.0931 | 0.0275 | -7.99 | 1 | 34 | 0.00001626 | deleterious |
| R121H | R | 121 | H | 0.4219 | 0.3415 | -4.99 | 1 | 34 | 0.00003252 | deleterious |
| R121L | R | 121 | L | 0.1677 | 0.172 | -6.99 | 1 | 34 | NA | deleterious |
| R121S | R | 121 | S | 0.0329 | 0.0929 | -5.99 | 1 | 32 | NA | deleterious |
| R125Q | R | 125 | Q | 0.9589 | 0.9318 | -3.9 | 0.999 | 35 | 1.45E-05 | deleterious |
| R125W | R | 125 | W | 0.5663 | 0.9159 | -7.89 | 1 | 32 | NA | deleterious |
| M126V | M | 126 | V | 0.191 | 0.2141 | -4 | 0.997 | 25.2 | NA | deleterious |
| R132S | R | 132 | S | 0.8995 | 0.9526 | -3.18 | 0.465 | 20.3 | 1.22E-05 | neutral |
| R132C | R | 132 | C | 0.9934 | 0.9579 | -5.1 | 0.997 | 27.1 | 2.86E-04 | neutral |
| R132H | R | 132 | H | 0.6791 | 0.697 | -3.25 | 0.994 | 26 | 8.14E-06 | neutral |
| D133N | D | 133 | N | 0.7145 | 0.7911 | -1 | 0.002 | 18.75 | 1.87E-04 | neutral |
| G134R | G | 134 | R | 0.8359 | 0.788 | -7.96 | 1 | 27.4 | 8.68E-05 | neutral |
| T135M | T | 135 | M | 0.7668 | 0.9235 | 0.54 | 0.036 | 2.674 | 4.70E-05 | neutral |
| T141M | T | 141 | M | 0.9102 | 0.9566 | -4.83 | 1 | 26.4 | 1.22E-05 | neutral |
| E144K | E | 144 | K | 0.1526 | 0.1619 | -4 | 1 | 27.1 | 3.26E-05 | deleterious |
| P145L | P | 145 | L | 0.0032 | 0.0157 | -9.67 | 1 | 28.3 | 4.084E-06 | deleterious |
| G148R | G | 148 | R | 0.4629 | 0.4145 | -8 | 1 | 29.3 | 0.00001227 | deleterious |
| G151R | G | 151 | R | 0.0211 | 0.1515 | -8 | 1 | 28.9 | 0.00002548 | deleterious |
| G153R | G | 153 | R | 0.2129 | 0.1491 | -7.87 | 1 | 33 | NA | deleterious |
| A155V | A | 155 | V | 0.3387 | 0.4702 | -4 | 1 | 28.5 | NA | deleterious |
| A157P | A | 157 | P | 0.2766 | 0.2592 | -3.66 | 0.987 | 24.3 | NA | deleterious |
| C165Y | C | 165 | Y | 0.0862 | 0.1521 | -11 | 1 | 31 | NA | deleterious |
| V168A | V | 168 | A | 0.9309 | 0.7433 | -3.6 | 0.849 | 25.8 | NA | deleterious |
| V168M | V | 168 | M | 0.6908 | 0.5382 | -2.8 | 0.997 | 33 | NA | deleterious |
| E176K | E | 176 | K | 0.1263 | 0.4711 | -3.88 | 1 | 28.5 | NA | deleterious |
| V178M | V | 178 | M | 0.5845 | 0.6448 | -2.95 | 0.992 | 27 | 2.06E-05 | neutral |
| V180M | V | 180 | M | 0.5119 | 0.6866 | -2.16 | 0.969 | 27.1 | 2.05E-05 | neutral |
| V180A | V | 180 | A | 0.5269 | 0.7651 | -2.89 | 0.054 | 22.8 | NA | deleterious |
| R182W | R | 182 | W | 0.2397 | 0.6237 | -6.29 | 0.999 | 26.9 | 8.21E-06 | neutral |
| R182Q | R | 182 | Q | 0.5751 | 0.9804 | -2.33 | 0.304 | 23.8 | 2.05E-05 | neutral |
| A183V | A | 183 | V | 0.012 | 0.4579 | -3.73 | 0.859 | 26.5 | 1.23E-05 | neutral |
| T191M | T | 191 | M | 0.037 | 0.0529 | -5.69 | 1 | 27.5 | 0.00006148 | deleterious |
| R196S | R | 196 | S | 0.6447 | 0.709 | -0.17 | 0.002 | 12.38 | 3.63E-05 | neutral |
| D198V | D | 198 | V | 0.137 | 0.4332 | -8.49 | 0.993 | 27 | NA | deleterious |
| P200L | P | 200 | L | 0.7942 | 0.9659 | -9.25 | 0.992 | 27 | 1.64E-05 | deleterious |
| V204M | V | 204 | M | 0.5132 | 0.624 | -1.83 | 0.089 | 24.5 | 4.72E-05 | neutral |
| V206M | V | 206 | M | 0.9462 | 0.9529 | -2.61 | 0.968 | 29.8 | 1.82E-05 | neutral |
| R209W | R | 209 | W | 0.2941 | 0.5564 | -6.74 | 0.999 | 33 | 8.18E-06 | neutral |
| N212K | N | 212 | K | 0.8045 | 0.9781 | 0.78 | 0.001 | 3.6 | 1.18E-04 | neutral |

| | | | | | | | | | | |
|---|---|---|---|---|---|---|---|---|---|---|
| E213K | E | 213 | K | 0.6843 | 0.8792 | -3.3 | 0.286 | 24.8 | 2.04E-05 | neutral |
| P215S | P | 215 | S | 0.2989 | 0.7447 | -6.76 | 0.984 | 27 | 1.63E-05 | neutral |
| S217F | S | 217 | F | 0.0069 | 0.0269 | -5.17 | 0.999 | 33 | NA | deleterious |
| R224C | R | 224 | C | 0.384 | 0.8956 | -4.8 | 0.443 | 27.5 | 1.99E-04 | neutral |
| R224H | R | 224 | H | 0.3182 | 0.8545 | -3.3 | 0.295 | 32 | 8.124E-06 | deleterious |
| A226T | A | 226 | T | 0.1095 | 0.5922 | -1.1 | 0.1 | 24 | NA | deleterious |
| N228K | N | 228 | K | 0.2257 | 0.3521 | -6 | 1 | 28.7 | 8.12E-06 | deleterious |
| N228S | N | 228 | S | 0.1237 | 0.2577 | -5 | 1 | 26.1 | NA | deleterious |
| D234N | D | 234 | N | 0.7173 | 0.9493 | -4.91 | 0.994 | 28.4 | 0.00001083 | deleterious |
| A237T | A | 237 | T | 0.7773 | 0.9604 | -3.83 | 1 | 31 | 8.12E-06 | neutral |
| M250L | M | 250 | L | 0.7082 | 0.796 | -2.58 | 0.258 | 29.1 | 1.10E-05 | neutral |
| T257M | T | 257 | M | 0.0034 | 0.1657 | -6 | 1 | 26.3 | 0.00004735 | deleterious |
| G259S | G | 259 | S | 0.0289 | 0.0386 | -6 | 1 | 33 | 7.272E-06 | deleterious |
| T262M | T | 262 | M | 0.003 | 0.2648 | -4.94 | 1 | 33 | 0.00002043 | deleterious |
| T262R | T | 262 | R | 0.0051 | 0.009 | -4.97 | 1 | 32 | NA | deleterious |
| I264T | I | 264 | T | 0.6301 | 0.8783 | -4.2 | 0.591 | 26.5 | 1.09E-05 | neutral |
| R266K | R | 266 | K | 0.7266 | 0.9969 | -2.6 | 0.587 | 26.9 | 8.17E-06 | deleterious |
| R266G | R | 266 | G | 0.0628 | 0.6182 | -6.8 | 0.998 | 27.6 | NA | deleterious |
| C275Y | C | 275 | Y | 0.0483 | 0.0809 | -5.14 | 0.999 | 24.8 | NA | deleterious |
| I278T | I | 278 | T | 0.3088 | 0.3613 | -3.82 | 0.967 | 23.5 | 7.80E-04 | deleterious |
| D281N | D | 281 | N | 0.0198 | 0.0682 | -4.71 | 0.996 | 27.2 | NA | deleterious |
| E283K | E | 283 | K | 0.4311 | 0.8072 | -1.84 | 0.045 | 22.1 | 2.02E-05 | neutral |
| I286V | I | 286 | V | 0.9116 | 0.8638 | -0.65 | 0.015 | 9.536 | 1.13E-04 | neutral |
| A288T | A | 288 | T | 0.0164 | 0.1446 | -3.6 | 0.985 | 28 | NA | deleterious |
| E292V | E | 292 | V | 0.9118 | 0.9458 | -4.83 | 0.056 | 23.2 | 1.67E-05 | neutral |
| T296R | T | 296 | R | 0.4119 | 0.6774 | -3.7 | 0.09 | 14.4 | 7.94E-05 | neutral |
| T296M | T | 296 | M | 0.7583 | 0.9676 | -3.91 | 0.969 | 24.3 | 2.44E-04 | neutral |
| E302K | E | 302 | K | 0.2196 | 0.6586 | -2.29 | 0.195 | 23 | 4.06E-06 | deleterious |
| G305R | G | 305 | R | 0.0498 | 0.1637 | -7.85 | 1 | 31 | NA | deleterious |
| G307S | G | 307 | S | 0.3083 | 0.1277 | -5.62 | 1 | 33 | 1.59E-04 | deleterious |
| D309N | D | 309 | N | 0.1807 | 0.2649 | -4.54 | 0.996 | 33 | 1.63E-05 | neutral |
| T313M | T | 313 | M | 0.2751 | 0.6047 | -4.94 | 1 | 31 | 5.77E-05 | neutral |
| V314M | V | 314 | M | 0.0092 | 0.2303 | -2.78 | 1 | 32 | 1.22E-05 | neutral |
| T318M | T | 318 | M | 0.8941 | 0.9743 | -2.31 | 0.976 | 23 | 4.47E-05 | neutral |
| V320A | V | 320 | A | 0.2581 | 0.6943 | -3.92 | 0.926 | 25.9 | 1.63E-05 | deleterious |
| D321V | D | 321 | V | 0.5304 | 0.9307 | -8.83 | 1 | 25.9 | NA | deleterious |
| A331E | A | 331 | E | 0.3362 | 0.327 | -1.63 | 0.671 | 26.4 | 4.07E-06 | deleterious |
| A331V | A | 331 | V | 0.2468 | 0.7228 | -2.9 | 0.819 | 24.1 | 8.135E-06 | deleterious |
| R336C | R | 336 | C | 0.0127 | 0.1427 | -7.85 | 1 | 33 | 0.00001628 | deleterious |
| R336H | R | 336 | H | 0.05 | 0.3173 | -4.91 | 1 | 32 | NA | deleterious |

| | | | | | | | | | | |
|---|---|---|---|---|---|---|---|---|---|---|
| M337R | M | 337 | R | 0.8061 | 0.84 | 2.88 | 0.001 | 6.771 | 1.63E-05 | neutral |
| M337L | M | 337 | L | 0.6844 | 0.8067 | -1.17 | 0.001 | 22.7 | 8.14E-06 | neutral |
| L338P | L | 338 | P | 0.2793 | 0.4886 | -6.68 | 1 | 25.9 | NA | deleterious |
| A340V | A | 340 | V | 0.9653 | 0.8439 | -2.04 | 0.174 | 22.6 | 1.22E-05 | neutral |
| A340T | A | 340 | T | 0.1765 | 0.6132 | -0.47 | 0.009 | 8.364 | 8.15E-06 | neutral |
| G347S | G | 347 | S | 0.0468 | 0.1128 | -5.82 | 0.999 | 33 | 3.24E-05 | deleterious |
| S349N | S | 349 | N | 0.0261 | 0.1219 | -2.93 | 0.991 | 31 | NA | deleterious |
| A350S | A | 350 | S | 0.6903 | 0.8985 | 1.87 | 0 | 0.758 | 8.61E-06 | neutral |
| S352N | S | 352 | N | 0.0783 | 0.533 | -2.15 | 0.427 | 25.5 | NA | deleterious |
| T353M | T | 353 | M | 0.1632 | 0.9789 | -1.21 | 0.059 | 24.2 | 0.00003025 | deleterious |
| V354M | V | 354 | M | 0.5036 | 0.79 | 1.93 | 0.002 | 0.002 | NA | deleterious |
| A357G | A | 357 | G | 0.8994 | 0.9493 | -3.91 | 0.631 | 25.2 | 8.54E-06 | neutral |
| V358M | V | 358 | M | 0.906 | 0.8892 | -1.14 | 0.765 | 22 | 1.01E-04 | neutral |
| A361T | A | 361 | T | 0.7819 | 0.8319 | -3.7 | 0.61 | 24.3 | 1.86E-05 | deleterious |
| A361V | A | 361 | V | 0.2529 | 0.3578 | -3.61 | 0.993 | 25.1 | NA | deleterious |
| E366K | E | 366 | K | 0.8913 | 0.8827 | -2.14 | 0.003 | 22.7 | 8.31E-06 | neutral |
| R369H | R | 369 | H | 0.7695 | 0.8545 | -3.67 | 1 | 34 | 0.00003233 | deleterious |
| C370Y | C | 370 | Y | 0.0039 | 0.0156 | -9.84 | 1 | 31 | 0.00001236 | deleterious |
| V371M | V | 371 | M | 0.6711 | 0.8158 | -2.93 | 1 | 34 | NA | deleterious |
| V372I | V | 372 | I | 0.732 | 0.9047 | -0.97 | 0.204 | 28.5 | 1.23E-05 | neutral |
| D376N | D | 376 | N | 0.0267 | 0.1392 | -4.89 | 1 | 35 | 4.09E-06 | deleterious |
| R379Q | R | 379 | Q | 0.0644 | 0.0618 | -3.75 | 0.708 | 34 | 8.18E-06 | deleterious |
| K384N | K | 384 | N | 0.2378 | 0.157 | -4.67 | 0.999 | 27.4 | NA | deleterious |
| D388Y | D | 388 | Y | 0.1047 | 0.3063 | -8.56 | 0.989 | 32 | 1.22E-05 | neutral |
| M391I | M | 391 | I | 0.0167 | 0.0884 | -3.82 | 0.293 | 25.1 | NA | deleterious |

### Supplementary Table 4. Patient clinical phenotypes and vitamin $B_6$ responsiveness, and inferred enzyme activity from VE maps.

| Patient | Inferred amino acid change | Nucleotide change | Vitamin $B_6$ responsivity | Disease severity | Age of symptoms onset | Vascular | Connective tissue | Neuropsychiatric | Present age of patient | VE map inferred total CBS activity | | |
|---|---|---|---|---|---|---|---|---|---|---|---|---|
| | | | | | | | | | | low vitamin B6 | high vitamin B6 | high-low vitamin B6 |
| 1 | p.[A71Yfs*25];[V10Wfs*71] | c.[210-1G>C];[28delG] | 1 | 1 | 1 | 0 | 1 | 1 | 3 | 0.0000 | 0.0000 | 0.0000 |
| 2 | p.[I278T];[W409_G453del] | c.[833T>C];[1224-2A>C] | 2 | 3 | 6 | 0 | 1 | 0 | 13 | 0.0474 | 0.0600 | 0.0126 |
| 3 | p.[I278T];[E144K;A155T] | c.[833T>C];[430G>A;463G>A] | 2 | 2 | 2 | 0 | 1 | 0 | 10 | 0.0665 | 0.0805 | 0.0140 |
| 4 | p.[I278T];[E144K;A155T] | c.[833T>C];[430G>A;463G>A] | 2 | 2 | 6 | 0 | 1 | 0 | 6 | 0.0665 | 0.0805 | 0.0140 |
| 5 | p.[G246Dfs*52];[W409_G453del] | c.[828+1G>A];[1224-2A>C] | 1 | 1 | 1.5 | 0 | 1 | 1 | 10 | 0.0000 | 0.0000 | 0.0000 |
| 6 | p.[W409_G453del];[E144K;A155T] | c.[1224-2A>C];[430G>A;463G>A] | 1 | 3 | 7 | 0 | 1 | 0 | 9 | 0.0191 | 0.0205 | 0.0014 |
| 7 | p.[I278T];[I278T] | c.[833T>C];[833T>C] | 3 | 3 | 10 | 0 | 1 | 0 | 47 | 0.0948 | 0.1200 | 0.0252 |
| 8 | p.[C165Y];[C165Y] | c.[494G>A];[494G>A] | 1 | 1 | 4 | 0 | 1 | 1 | 5 | 0.0200 | 0.0381 | 0.0180 |
| 9 | p.[I278T];[A71Yfs*25] | c.[833T>C];[210-1G>C] | 1 | 1 | 27 | 1 | 1 | 1 | 29 | 0.0474 | 0.0600 | 0.0126 |
| 10 | p.[W409_G453del];[W409_G453del] | c.[1224-2A>C];[1224-2A>C] | 1 | 1 | 1 | 0 | 1 | 1 | 3 | 0.0000 | 0.0000 | 0.0000 |
| 11 | p.[I278T];[I278T] | c.[833T>C];[833T>C] | 4 | 4 | 29 | 1 | 0 | 0 | 32 | 0.0948 | 0.1200 | 0.0252 |
| 12 | p.[E144K;A155T];[?] | c.[430G>A;463G>A];[?] | 1 | 1 | 3 | 1 | 1 | 1 | 17 | 0.0191 | 0.0205 | 0.0014 |
| 13 | p.[I278T];[E144K;A155T] | c.[833T>C];[430G>A;463G>A] | 2 | 2 | 18 | 1 | 1 | 0 | 19 | 0.0665 | 0.0805 | 0.0140 |
| 14 | p.[I278T];[D376N] | c.[833T>C];[1126G>A] | 2 | 3 | 8 | 0 | 1 | 0 | 10 | 0.0503 | 0.0772 | 0.0269 |
| 15 | p.[I278T];[I278T] | c.[833T>C];[833T>C] | 3 | 3 | 21 | 1 | 1 | 0 | 30 | 0.0948 | 0.1200 | 0.0252 |
| 16 | p.[A71Yfs*25];[A71Yfs*25] | c.[210-1G>C];[210-1G>C] | 1 | 1 | 0.5 | 0 | 0 | 1 | 1.25 | 0.0000 | 0.0000 | 0.0000 |
| 17 | p.[A71Yfs*25];[A71Yfs*25] | c.[210-1G>C];[210-1G>C] | 1 | 1 | 5 | 0 | 1 | 1 | 9 | 0.0000 | 0.0000 | 0.0000 |
| 18 | p.[I278T];[I278T] | c.[833T>C];[833T>C] | 3 | 4 | 21 | 1 | 0 | 0 | 35 | 0.0948 | 0.1200 | 0.0252 |
| 19 | p.[P49L(;)R336H] | c.[146C>T(;)1007G>A] | 4 | 4 | 40 | 1 | 0 | 0 | 54 | 0.0283 | 0.1101 | 0.0818 |
| 20 | p.[P49L(;)R336H] | c.[146C>T(;)1007G>A] | 4 | 3 | 46 | 1 | 1 | 0 | 51 | 0.0283 | 0.1101 | 0.0818 |
| 21 | p.[P49L(;)R336H] | c.[146C>T(;)1007G>A] | 4 | 5 | NA (asymtpomatic, detected by family screening) | 0 | 0 | 0 | 47 | 0.0283 | 0.1101 | 0.0818 |
| 22 | p.[I278T];[I278T] | c.[833T>C];[833T>C] | 3 | 4 | 19 | 1 | 0 | 0 | 44 | 0.0948 | 0.1200 | 0.0252 |
| 23 | p.[I278T];[I278T] | c.[833T>C];[833T>C] | 3 | 3 | 22 | 0 | 1 | 0 | 44 | 0.0948 | 0.1200 | 0.0252 |
| 24 | p.[I278T];[R336C] | c.[833T>C];[1006C>T] | 3 | 3 | 24 | 1 | 1 | 0 | 73 | 0.0488 | 0.0777 | 0.0289 |
| 25 | p.[P145L(;)I278T] | c.[434C>T(;)833T>C] | 3 | 3 | 16 | 0 | 1 | 0 | 27 | 0.0477 | 0.0617 | 0.0140 |
| 26 | p.[Q7Pfs*30];[E144K;A155T] | c.[19dupC];[430G>A,463G>A] | 1 | NA | NA (newborn screening) | 0 | 0 | 0 | 0.025 | 0.0191 | 0.0205 | 0.0014 |
| 27 | p.[A71Yfs*25(;)K211*] | c.[210-1G>C(;)631A>T] | 1 | 1 | 5 | 1 | 1 | 1 | 33 | 0.0000 | 0.0000 | 0.0000 |
| 28 | p.[W409_G453del];[W409_G453del] | c.[1224-2A>C];[1224-2A>C] | 1 | NA | NA (newborn screening) | 0 | 0 | 0 | 0.025 | 0.0000 | 0.0000 | 0.0000 |

Vitamin $B_6$ responsivity based on response to vitamin $B_6$ administration and necessity of other treatments: 1 nonresponsive (needs diet and betaine, may be receiving vitamin $B_6$ due to historical reasons), 2 partial responsive (needs large doses of pyridoxine and varying degree of low protein diet), 3 fully responsive (needs only pyridoxine, doses > 0.5 mg/kg/day) and 4 extremely pyridoxine responsive (needs only doses < 0.5 mg/kg/day of pyridoxine per day) to achieve good therapeutic control (tHcy < 50 umol/L)

Severity of disease- takes into account which organ systems were affected by the disease (may include also symptoms manifesting between the first symptoms and the time of diagnosis, or complication occurring in poorly compliant patients); severity cannot be determined in patients detected by newborn screening as they are treated since neonatal period. 5-no symptoms at time of diagnosis (in a patient detected by family screening); 4-mild disease (only thrombosis in any vascular bed with no other symptoms); 3-moderate disease (connective tissue involvement with or without thrombosis); 2- borderline severity mild cognitive impairment with good social outcome, regardless of other somatic complications) 1- severe disease (severe neuropsychiatric complications including poor social outcome, regardless of other somatic complications)

Supplementary Table 5. List of enzymes containing cofactors.

| Enzyme name | Cofactor | UniprotID | Enzyme name | official gene name | Disease associated |
|---|---|---|---|---|---|
| Glutathione-disulfide reductase | FAD | P00390 | GSHR | *GSR* | yes |
| L-amino-acid oxidase | FAD | Q96RQ9 | OXLA | *IL4I1* | |
| Fructose-bisphosphate aldolase | Zn(2+) | P09972 | ALDOC | *ALDOC* | |
| Protein-glutamine gamma-glutamyltransferase | Ca(2+) | P49221 | TGM4 | *TGM4* | |
| Protein-glutamine gamma-glutamyltransferase | Ca(2+) | Q96PF1 | TGM7 | *TGM7* | |
| Xaa-Pro aminopeptidase | Cobalt cation or Mn(2+) | Q9NQH7 | XPP3 | *XPNPEP3* | yes |
| Selenocysteine lyase | Pyridoxal 5'-phosphate | Q96I15 | SCLY | *SCLY* | |
| Sodium/potassium-exchanging ATPase | Mg(2+) | Q13733 | AT1A4 | *ATP1A4* | |
| Long-chain-acyl-CoA dehydrogenase | FAD | P28330 | ACADL | *ACADL* | yes |
| Carbonic anhydrase | Zn(2+) | P22748 | CAH4 | *CA4* | yes |
| Unspecific monooxygenase | Heme-thiolate | Q9HB55 | CP343 | *CYP3A43* | |
| Prostaglandin-D synthase | Glutathione | P41222 | PTGDS | *PTGDS* | |
| Phospholipid-translocating ATPase | Mg(2+) | Q8TF62 | AT8B4 | *ATP8B4* | |
| Protein-glutamine gamma-glutamyltransferase | Ca(2+) | Q08188 | TGM3 | *TGM3* | |
| Glutathione peroxidase | Se(2+) | Q8TED1 | GPX8 | *GPX8* | |
| Carbonic anhydrase | Zn(2+) | Q9Y2D0 | CAH5B | *CA5B* | |
| Phospholipid-translocating ATPase | Mg(2+) | Q9P241 | AT10D | *ATP10D* | |
| Ste24 endopeptidase | Zn(2+) | O75844 | FACE1 | *ZMPSTE24* | yes |
| Cholesterol 25-hydroxylase | Fe cation | O95992 | CH25H | *CH25H* | |
| Methylcrotonoyl-CoA carboxylase | Biotin | Q96RQ3 | MCCA | *MCCC1* | yes |
| N-acetylgalactosamine kinase | Mg(2+) | Q01415 | GALK2 | *GALK2* | |
| UDP-sugar diphosphatase | Divalent cation | O95848 | NUD14 | *NUDT14* | |
| Apyrase | Ca(2+) | O75355 | ENTP3 | *ENTPD3* | |
| Methionyl aminopeptidase | Cobalt cation | Q6UB28 | MAP12 | *METAP1D* | |
| 24-hydroxycholesterol 7-alpha-hydroxylase | Heme-thiolate | Q9NYL5 | CP39A | *CYP39A1* | |
| Alcohol dehydrogenase (NADP(+)) | Zn(2+) | P14550 | AK1A1 | *AKR1A1* | |
| Unspecific monooxygenase | Heme-thiolate | P33260 | CP2CI | *CYP2C18* | |
| Trimethylamine monooxygenase | Flavoprotein | P31513 | FMO3 | *FMO3* | yes |
| Alkylglycerol monooxygenase | Glutathione | Q6ZNB7 | ALKMO | *AGMO* | |
| Interstitial collagenase | Zn(2+) | P03956 | MMP1 | *MMP1* | yes |
| Calpain-2 | Ca(2+) | P17655 | CAN2 | *CAPN2* | |
| Prostaglandin-D synthase | Glutathione | O60760 | HPGDS | *HPGDS* | |
| Xylosylprotein 4-beta-galactosyltransferase | Mn(2+) | Q9UBV7 | B4GT7 | *B4GALT7* | yes |
| Sulfinoalanine decarboxylase | Pyridoxal 5'-phosphate | Q9Y600 | CSAD | *CSAD* | |
| Hyaluronan synthase | Mg(2+) | Q92839 | HYAS1 | *HAS1* | |
| Protein-glutamine gamma-glutamyltransferase | Ca(2+) | O95932 | TGM3L | *TGM6* | yes |
| Acetyl-CoA carboxylase | Biotin | O00763 | ACACB | *ACACB* | |

| | | | | | |
|---|---|---|---|---|---|
| Kynurenine--oxoglutarate transaminase | Pyridoxal 5'-phosphate | P00505 | AATM | *GOT2* | |
| tRNA(Ala)(adenine(37)) deaminase | Mg(2+) | Q9BUB4 | ADAT1 | *ADAT1* | |
| Pyridoxal phosphatase | Mg(2+) | Q96GD0 | PLPP | *PDXP* | |
| Threonine ammonia-lyase | Pyridoxal 5'-phosphate or iron-sulfur | Q96GA7 | SDSL | *SDSL* | |
| Protein geranylgeranyltransferase type I | Zn(2+) | P49354 | FNTA | *FNTA* | |
| 7-alpha-hydroxycholest-4-en-3-one 12-alpha-hydroxylase | Heme-thiolate | Q9UNU6 | CP8B1 | *CYP8B1* | |
| Inositol-trisphosphate 3-kinase | Ca(2+) | Q96DU7 | IP3KC | *ITPKC* | yes |
| Serine racemase | Pyridoxal 5'-phosphate | Q9GZT4 | SRR | *SRR* | |
| Protein-glutamine gamma-glutamyltransferase | Ca(2+) | O43548 | TGM5 | *TGM5* | yes |
| ADAM10 endopeptidase | Zn(2+) | O14672 | ADA10 | *ADAM10* | yes |
| Adrenodoxin-NADP(+) reductase | FAD | P22570 | ADRO | *FDXR* | |
| Procollagen N-endopeptidase | Zn(2+) | O95450 | ATS2 | *ADAMTS2* | yes |
| Galactosylgalactosylxylosylprotein 3-beta-glucuronosyltransferase | Mn(2+) | Q9P2W7 | B3GA1 | *B3GAT1* | |
| Phosphopyruvate hydratase | Mg(2+) | P06733 | ENOA | *ENO1* | |
| Phospholipase A(2) | Ca(2+) | P04054 | PA21B | *PLA2G1B* | |
| [Myosin light-chain] kinase | Ca(2+) | Q32MK0 | MYLK3 | *MYLK3* | |
| Sulfinoalanine decarboxylase | Pyridoxal 5'-phosphate | Q6ZQY3 | GADL1 | *GADL1* | |
| [Histone H3]-lysine-36 demethylase | Fe(2+) | Q9Y2K7 | KDM2A | *KDM2A* | |
| Acetyl-CoA carboxylase | Biotin | Q13085 | ACACA | *ACACA* | yes |
| UDP-sugar diphosphatase | Divalent cation | Q9BRQ3 | NUD22 | *NUDT22* | |
| (S)-2-hydroxy-acid oxidase | FMN | Q9UJM8 | HAOX1 | *HAO1* | |
| Alcohol dehydrogenase | Zn(2+) or Fe cation | P08319 | ADH4 | *ADH4* | |
| Prenylcysteine oxidase | FAD | Q9UHG3 | PCYOX | *PCYOX1* | |
| Dopamine beta-monooxygenase | Cu cation | P09172 | DOPO | *DBH* | yes |
| Cholestanetriol 26-monooxygenase | Heme-thiolate | Q02318 | CP27A | *CYP27A1* | yes |
| Prostaglandin-E synthase | Glutathione | Q15185 | TEBP | *PTGES3* | |
| Calcium/calmodulin-dependent protein kinase | Ca(2+) | Q13554 | KCC2B | *CAMK2B* | |
| Leukotriene-B(4) 20-monooxygenase | Heme-thiolate | Q08477 | CP4F3 | *CYP4F3* | |
| Inositol-trisphosphate 3-kinase | Ca(2+) | P23677 | IP3KA | *ITPKA* | |
| Cysteamine dioxygenase | Fe cation | Q96SZ5 | AEDO | *ADO* | |
| Serine C-palmitoyltransferase | Pyridoxal 5'-phosphate | Q9NUV7 | SPTC3 | *SPTLC3* | |
| Hyaluronan synthase | Mg(2+) | Q92819 | HYAS2 | *HAS2* | |
| Sterol 14-alpha-demethylase | Heme-thiolate | Q16850 | CP51A | *CYP51A1* | yes |
| Phospholipase A(2) | Ca(2+) | Q9NZK7 | PA2GE | *PLA2G2E* | |
| Dimethylglycine dehydrogenase | FAD | Q9UI17 | M2GD | *DMGDH* | yes |
| Acyl-CoA 6-desaturase | Fe cation | O95864 | FADS2 | *FADS2* | |
| Glutamate carboxypeptidase II | Zn(2+) | Q9UQQ1 | NALDL | *NAALADL1* | |
| (S)-2-hydroxy-acid oxidase | FMN | Q9NYQ3 | HAOX2 | *HAO2* | |
| Membrane dipeptidase | Zn(2+) | Q9H4B8 | DPEP3 | *DPEP3* | |
| Peptidylglycine monooxygenase | Cu cation | P19021 | AMD | *PAM* | yes |

| | | | | | |
|---|---|---|---|---|---|
| Tryptophan 2,3-dioxygenase | Heme | P48775 | T23O | *TDO2* | |
| Carbonic anhydrase | Zn(2+) | P43166 | CAH7 | *CA7* | |
| Tyrosine transaminase | Pyridoxal 5'-phosphate | P17735 | ATTY | *TAT* | yes |
| Alcohol dehydrogenase | Zn(2+) or Fe cation | P07327 | ADH1A | *ADH1A* | |
| Unspecific monooxygenase | Heme-thiolate | Q7Z449 | CP2U1 | *CYP2U1* | yes |
| Serine C-palmitoyltransferase | Pyridoxal 5'-phosphate | O15269 | SPTC1 | *SPTLC1* | yes |
| Squalene synthase | Mn(2+) or Mg(2+) | P37268 | FDFT | *FDFT1* | |
| Arachidonate 15-lipoxygenase | Fe cation | P16050 | LOX15 | *ALOX15* | |
| Glutamate decarboxylase | Pyridoxal 5'-phosphate | Q05329 | DCE2 | *GAD2* | |
| Mannose-6-phosphate isomerase | Zn(2+) | P34949 | MPI | *MPI* | yes |
| Phospholipid-translocating ATPase | Mg(2+) | O60312 | AT10A | *ATP10A* | yes |
| Dihydroorotate dehydrogenase (quinone) | FMN | Q02127 | PYRD | *DHODH* | yes |
| Flavin-containing monooxygenase | FAD | O60774 | FMO6 | *FMO6P* | |
| Thromboxane-A synthase | Heme-thiolate | P24557 | THAS | *TBXAS1* | yes |
| Lysine carboxypeptidase | Zn(2+) | P15169 | CBPN | *CPN1* | yes |
| Sodium/potassium-exchanging ATPase | Mg(2+) | P05023 | AT1A1 | *ATP1A1* | yes |
| Ornithine aminotransferase | Pyridoxal 5'-phosphate | P04181 | OAT | *OAT* | yes |
| Alcohol dehydrogenase | Zn(2+) or Fe cation | P28332 | ADH6 | *ADH6* | |
| Tyrosinase | Cu cation | P14679 | TYRO | *TYR* | yes |
| Dihydrolipoyl dehydrogenase | FAD | P09622 | DLDH | *DLD* | yes |
| Calcium-transporting ATPase | Mg(2+) | Q16720 | AT2B3 | *ATP2B3* | yes |
| Membrane alanyl aminopeptidase | Zn(2+) | P15144 | AMPN | *ANPEP* | |
| Calcium-transporting ATPase | Mg(2+) | P23634 | AT2B4 | *ATP2B4* | |
| Unspecific monooxygenase | Heme-thiolate | Q96SQ9 | CP2S1 | *CYP2S1* | |
| Phospholipase A(1) | Ca(2+) | Q8WU67 | ABHD3 | *ABHD3* | |
| Carbonic anhydrase | Zn(2+) | Q8N1Q1 | CAH13 | *CA13* | |
| Phosphatidate phosphatase | Mg(2+) | Q9BQK8 | LPIN3 | *LPIN3* | yes |
| Procollagen C-endopeptidase | Zn(2+) | P13497 | BMP1 | *BMP1* | yes |
| Glutamate decarboxylase | Pyridoxal 5'-phosphate | Q99259 | DCE1 | *GAD1* | yes |
| Homogentisate 1,2-dioxygenase | Fe cation | Q93099 | HGD | *HGD* | yes |
| Phospholipid-translocating ATPase | Mg(2+) | Q9Y2Q0 | AT8A1 | *ATP8A1* | |
| Phospholipase A(2) | Ca(2+) | Q9BZM2 | PA2GF | *PLA2G2F* | |
| 17-alpha-hydroxyprogesterone deacetylase | Heme-thiolate | P05093 | CP17A | *CYP17A1* | yes |
| Phospholipase A(2) | Ca(2+) | Q3MJ16 | PA24E | *PLA2G4E* | |
| Endothelin-converting enzyme 1 | Zn(2+) | P42892 | ECE1 | *ECE1* | yes |
| Isocitrate dehydrogenase (NAD(+)) | Mn(2+) or Mg(2+) | P50213 | IDH3A | *IDH3A* | |
| Calcium-transporting ATPase | Mg(2+) | P98194 | AT2C1 | *ATP2C1* | yes |
| Pyroglutamyl-peptidase II | Zn(2+) | Q9UKU6 | TRHDE | *TRHDE* | |
| N-acyl-aliphatic-L-amino acid amidohydrolase | Zn(2+) | Q03154 | ACY1 | *ACY1* | yes |
| Carboxypeptidase E | Zn(2+) | P16870 | CBPE | *CPE* | |

| | | | | | |
|---|---|---|---|---|---|
| Pseudouridine 5'-phosphatase | Mg(2+) | Q08623 | HDHD1 | *HDHD1* | |
| Cytidine deaminase | Zn(2+) | P32320 | CDD | *CDA* | |
| Globotriaosylceramide 3-beta-N-acetylgalactosaminyltransferase | Mn(2+) | O75752 | B3GL1 | *B3GALNT1* | yes |
| 3-hydroxyanthranilate 3,4-dioxygenase | Fe cation | P46952 | 3HAO | *HAAO* | |
| Pyruvate dehydrogenase (acetyl-transferring) | Thiamine diphosphate | P11177 | ODPB | *PDHB* | yes |
| Cytochrome-b5 reductase | FAD | Q9UHQ9 | NB5R1 | *CYB5R1* | |
| Methylcytosine dioxygenase | Fe(2+) | O43151 | TET3 | *TET3* | |
| Calcium/calmodulin-dependent protein kinase | Ca(2+) | Q8IU85 | KCC1D | *CAMK1D* | |
| Calcium/calmodulin-dependent protein kinase | Ca(2+) | Q9UQM7 | KCC2A | *CAMK2A* | |
| Peroxidase | Heme | Q92626 | PXDN | *PXDN* | yes |
| Glycogenin glucosyltransferase | Mn(2+) | P46976 | GLYG | *GYG1* | yes |
| Calcium/calmodulin-dependent protein kinase | Ca(2+) | Q13557 | KCC2D | *CAMK2D* | |
| Phospholipase A(2) | Ca(2+) | Q86XP0 | PA24D | *PLA2G4D* | |
| Calcium/calmodulin-dependent protein kinase | Ca(2+) | Q8N5S9 | KKCC1 | *CAMKK1* | |
| Carboxypeptidase A | Zn(2+) | P15088 | CBPA3 | *CPA3* | |
| Phospholipase A(2) | Ca(2+) | O15496 | PA2GX | *PLA2G10* | |
| Cholesterol 24-hydroxylase | Heme-thiolate | Q9Y6A2 | CP46A | *CYP46A1* | |
| 8-oxo-dGTP diphosphatase | Mg(2+) | P36639 | 8ODP | *NUDT1* | |
| Selenide, water dikinase | Mg(2+) | P49903 | SPS1 | *SEPHS1* | |
| Steroid 21-monooxygenase | Heme-thiolate | P08686 | CP21A | *CYP21A2* | yes |
| Phospholipase A(2) | Ca(2+) | Q6P1J6 | PLB1 | *PLB1* | |
| Insulysin | Zn(2+) | P14735 | IDE | *IDE* | |
| Transketolase | Thiamine diphosphate | P29401 | TKT | *TKT* | |
| Mitochondrial processing peptidase | Zn(2+) | O75439 | MPPB | *PMPCB* | |
| Phospholipid-translocating ATPase | Mg(2+) | Q9NTI2 | AT8A2 | *ATP8A2* | yes |
| tRNA(Phe) (7-(3-amino-3-carboxypropyl)wyosine(37)-C(2))-hydroxylase | Fe(2+) | A2RUC4 | TYW5 | *TYW5* | |
| Hydroxyproline dehydrogenase | FAD | Q9UF12 | HYPDH | *PRODH2* | |
| Cysteine desulfurase | Pyridoxal 5'-phosphate | Q9Y697 | NFS1 | *NFS1* | |
| Alcohol dehydrogenase | Zn(2+) or Fe cation | P40394 | ADH7 | *ADH7* | |
| Phospholipid-translocating ATPase | Mg(2+) | Q8NB49 | AT11C | *ATP11C* | |
| Unspecific monooxygenase | Heme-thiolate | P20853 | CP2A7 | *CYP2A7* | |
| L-dopachrome isomerase | Zn(2+) | P40126 | TYRP2 | *DCT* | |
| 5-aminolevulinate synthase | Pyridoxal 5'-phosphate | P13196 | HEM1 | *ALAS1* | |
| Tryptophan 5-monooxygenase | Fe cation | Q8IWU9 | TPH2 | *TPH2* | yes |
| Phosphatidate phosphatase | Mg(2+) | Q5VZY2 | PLPP4 | *PPAPDC1A* | |
| Peptidyl-dipeptidase A | Zn(2+) | P12821 | ACE | *ACE* | yes |
| D-aminoacyl-tRNA deacylase | Zn(2+) | Q8TEA8 | DTD1 | *DTD1* | |
| Short-chain acyl-CoA dehydrogenase | FAD | P16219 | ACADS | *ACADS* | yes |
| Prostaglandin-E synthase | Glutathione | O14684 | PTGES | *PTGES* | |
| Very-long-chain acyl-CoA dehydrogenase | FAD | P49748 | ACADV | *ACADVL* | yes |

| | | | | | |
|---|---|---|---|---|---|
| Membrane-type matrix metalloproteinase-1 | Zn(2+) | P50281 | MMP14 | *MMP14* | yes |
| Monoamine oxidase | FAD | P27338 | AOFB | *MAOB* | |
| Cysteine-S-conjugate beta-lyase | Pyridoxal 5'-phosphate | Q16773 | KAT1 | *CCBL1* | |
| Ferroxidase | Cu cation | P00450 | CERU | *CP* | yes |
| Acireductone dioxygenase (Fe(2+)-requiring) | Fe(2+) | Q9BV57 | MTND | *ADI1* | |
| Sarcosine dehydrogenase | FMN | Q9UL12 | SARDH | *SARDH* | yes |
| Persulfide dioxygenase | Fe cation | O95571 | ETHE1 | *ETHE1* | yes |
| Inositol-3-phosphate synthase | NAD(+) | Q9NPH2 | INO1 | *ISYNA1* | |
| Phospholipid-translocating ATPase | Mg(2+) | O94823 | AT10B | *ATP10B* | |
| Phosphatidylinositol-3,4,5-trisphosphate 3-phosphatase | Mg(2+) | P60484 | PTEN | *PTEN* | |
| Unspecific monooxygenase | Heme-thiolate | Q86W10 | CP4Z1 | *CYP4Z1* | |
| Cystinyl aminopeptidase | Zn(2+) | Q9UIQ6 | LCAP | *LNPEP* | |
| Phospholipase A(2) | Ca(2+) | Q9UNK4 | PA2GD | *PLA2G2D* | |
| Flavin-containing monooxygenase | FAD | P49326 | FMO5 | *FMO5* | |
| N-acetylphosphatidylethanolamine-hydrolyzing phospholipase D | Zn(2+) | Q6IQ20 | NAPEP | *NAPEPLD* | |
| 6-pyruvoyltetrahydropterin synthase | Mg(2+) | Q03393 | PTPS | *PTS* | yes |
| Neurolysin | Zn(2+) | Q9BYT8 | NEUL | *NLN* | |
| ADAMTS-4 endopeptidase | Zn(2+) | O75173 | ATS4 | *ADAMTS4* | |
| Transketolase | Thiamine diphosphate | Q9H0I9 | TKTL2 | *TKTL2* | |
| Methylenetetrahydrofolate reductase (NAD(P)H) | FAD | P42898 | MTHFR | *MTHFR* | yes |
| Phosphorylase kinase | Ca(2+) | P15735 | PHKG2 | *PHKG2* | yes |
| Phospholipase A(2) | Ca(2+) | P39877 | PA2G5 | *PLA2G5* | yes |
| Xaa-Pro aminopeptidase | Cobalt cation or Mn(2+) | Q9NQW7 | XPP1 | *XPNPEP1* | |
| Aconitate hydratase | Iron-sulfur | Q99798 | ACON | *ACO2* | yes |
| Calcium-transporting ATPase | Mg(2+) | O75185 | AT2C2 | *ATP2C2* | |
| [Myosin light-chain] kinase | Ca(2+) | Q9H1R3 | MYLK2 | *MYLK2* | yes |
| Unspecific monooxygenase | Heme-thiolate | P05177 | CP1A2 | *CYP1A2* | yes |
| Glycerol-3-phosphate dehydrogenase | Flavin | P43304 | GPDM | *GPD2* | yes |
| D-aspartate oxidase | FAD | Q99489 | OXDD | *DDO* | |
| Arginase | Mn(2+) | P78540 | ARGI2 | *ARG2* | |
| Deoxyhypusine synthase | NAD(+) | P49366 | DHYS | *DHPS* | |
| Arachidonate 15-lipoxygenase | Fe cation | O15296 | LX15B | *ALOX15B* | |
| Retinal dehydrogenase | FAD | P00352 | AL1A1 | *ALDH1A1* | |
| NADPH:quinone reductase | Zn(2+) | Q08257 | QOR | *CRYZ* | |
| Glycine C-acetyltransferase | Pyridoxal 5'-phosphate | O75600 | KBL | *GCAT* | |
| Calcidiol 1-monooxygenase | Heme-thiolate | O15528 | CP27B | *CYP27B1* | yes |
| Phospholipase A(2) | Ca(2+) | Q9BZM1 | PG12A | *PLA2G12A* | |
| Propionyl-CoA carboxylase | Biotin | P05166 | PCCB | *PCCB* | yes |
| UDP-glucuronate decarboxylase | NAD(+) | Q8NBZ7 | UXS1 | *UXS1* | |
| Neprilysin | Zn(2+) | P08473 | NEP | *MME* | yes |

| | | | | | |
|---|---|---|---|---|---|
| [Elongation factor 2] kinase | Ca(2+) | O00418 | EF2K | *EEF2K* | |
| Protein geranylgeranyltransferase type I | Zn(2+) | P53609 | PGTB1 | *PGGT1B* | |
| Peptide deformylase | Fe(2+) | Q9HBH1 | DEFM | *PDF* | |
| Alanine--glyoxylate transaminase | Pyridoxal 5'-phosphate | Q9BYV1 | AGT2 | *AGXT2* | yes |
| Acyl-CoA oxidase | FAD | O15254 | ACOX3 | *ACOX3* | |
| [Myosin light-chain] kinase | Ca(2+) | Q15746 | MYLK | *MYLK* | yes |
| N-acyl-aromatic-L-amino acid amidohydrolase | Zn(2+) | Q96HD9 | ACY3 | *ACY3* | |
| Aromatic-L-amino-acid decarboxylase | Pyridoxal 5'-phosphate | P20711 | DDC | *DDC* | yes |
| Phospholipase A(1) | Ca(2+) | P53816 | PA216 | *PLA2G16* | |
| Phospholipase A(2) | Ca(2+) | Q9NZ20 | PA2G3 | *PLA2G3* | |
| Phosphatidate phosphatase | Mg(2+) | Q92539 | LPIN2 | *LPIN2* | yes |
| Ribonucleoside-diphosphate reductase | Fe(3+) or adenosylcob(III)alamin or Mn(2+) | Q7LG56 | RIR2B | *RRM2B* | yes |
| Cytochrome-b5 reductase | FAD | Q6BCY4 | NB5R2 | *CYB5R2* | |
| (R)-limonene 6-monooxygenase | Heme-thiolate | P33261 | CP2CJ | *CYP2C19* | yes |
| Phosphatidate phosphatase | Mg(2+) | Q14693 | LPIN1 | *LPIN1* | yes |
| Arginase | Mn(2+) | P05089 | ARGI1 | *ARG1* | yes |
| Protein-glutamine gamma-glutamyltransferase | Ca(2+) | P22735 | TGM1 | *TGM1* | yes |
| Ethanolamine-phosphate phospho-lyase | Pyridoxal 5'-phosphate | Q8TBG4 | AT2L1 | *ETNPPL* | |
| Flavin-containing monooxygenase | FAD | Q01740 | FMO1 | *FMO1* | |
| Coproporphyrinogen oxidase | Fe cation | P36551 | HEM6 | *CPOX* | yes |
| 2-(3-amino-3-carboxypropyl)histidine synthase | Iron-sulfur | Q9BQC3 | DPH2 | *DPH2* | |
| Carboxypeptidase U | Zn(2+) | Q96IY4 | CBPB2 | *CPB2* | |
| Isocitrate dehydrogenase (NADP(+)) | Mn(2+) or Mg(2+) | O75874 | IDHC | *IDH1* | yes |
| Pyridoxal 5'-phosphate synthase | FMN | Q9NVS9 | PNPO | *PNPO* | yes |
| Carbonic anhydrase | Zn(2+) | P35218 | CAH5A | *CA5A* | |
| Phospholipid-translocating ATPase | Mg(2+) | Q9Y2G3 | AT11B | *ATP11B* | |
| Isocitrate dehydrogenase (NADP(+)) | Mn(2+) or Mg(2+) | P48735 | IDHP | *IDH2* | yes |
| Tryptophan 5-monooxygenase | Fe cation | P17752 | TPH1 | *TPH1* | yes |
| Phosphopyruvate hydratase | Mg(2+) | P13929 | ENOB | *ENO3* | yes |
| 5-aminolevulinate synthase | Pyridoxal 5'-phosphate | P22557 | HEM0 | *ALAS2* | yes |
| Glycine dehydrogenase (aminomethyl-transferring) | Pyridoxal 5'-phosphate | P23378 | GCSP | *GLDC* | yes |
| D-amino-acid oxidase | FAD | P14920 | OXDA | *DAO* | yes |
| Iodide peroxidase | Heme | P07202 | PERT | *TPO* | yes |
| Mn(2+)-dependent ADP-ribose/CDP-alcohol diphosphatase | Mn(2+) | Q3LIE5 | ADPRM | *ADPRM* | |
| Phosphorylase kinase | Ca(2+) | Q16816 | PHKG1 | *PHKG1* | |
| Phospholipase A(2) | Ca(2+) | P47712 | PA24A | *PLA2G4A* | yes |
| Cytochrome-b5 reductase | FAD | Q7L1T6 | NB5R4 | *CYB5R4* | |
| Xaa-Pro aminopeptidase | Cobalt cation or Mn(2+) | O43895 | XPP2 | *XPNPEP2* | yes |
| Methionyl aminopeptidase | Cobalt cation | Q9HAU8 | RNPL1 | *RNPEPL1* | |

| | | | | | |
|---|---|---|---|---|---|
| Flavin-containing monooxygenase | FAD | Q99518 | FMO2 | *FMO2* | |
| Glutamyl aminopeptidase | Zn(2+) | Q07075 | AMPE | *ENPEP* | |
| Unspecific monooxygenase | Heme-thiolate | P04798 | CP1A1 | *CYP1A1* | |
| tRNA 4-demethylwyosine synthase (AdoMet-dependent) | Iron-sulfur | Q6NUM6 | TYW1B | *TYW1B* | |
| Carotenoid-9',10'-cleaving dioxygenase | Fe cation | Q9BYV7 | BCDO2 | *BCO2* | |
| Cardiolipin synthase (CMP-forming) | Divalent cation | Q9UJA2 | CRLS1 | *CRLS1* | |
| Transketolase | Thiamine diphosphate | P51854 | TKTL1 | *TKTL1* | |
| Membrane dipeptidase | Zn(2+) | Q9H4A9 | DPEP2 | *DPEP2* | |
| Glutamate formimidoyltransferase | Pyridoxal 5'-phosphate | O95954 | FTCD | *FTCD* | yes |
| Calpain-2 | Ca(2+) | A6NHC0 | CAN8 | *CAPN8* | |
| Monoamine oxidase | FAD | P21397 | AOFA | *MAOA* | yes |
| Fructose-bisphosphate aldolase | Zn(2+) | P05062 | ALDOB | *ALDOB* | yes |
| Ribonucleoside-diphosphate reductase | Fe(3+) or adenosylcob(III)alamin or Mn(2+) | P23921 | RIR1 | *RRM1* | |
| Alanine transaminase | Pyridoxal 5'-phosphate | Q8TD30 | ALAT2 | *GPT2* | |
| Proline dehydrogenase | FAD | O43272 | PROD | *PRODH* | yes |
| Endothelin-converting enzyme 1 | Zn(2+) | P0DPD8 | EFCE2 | *EEF1AKMT4-ECE2* | |
| Ornithine decarboxylase | Pyridoxal 5'-phosphate | P11926 | DCOR | *ODC1* | yes |
| Molybdopterin molybdotransferase | Zn(2+) or Mg(2+) | Q9NQX3 | GEPH | *GPHN* | yes |
| Phosphopantothenoylcysteine decarboxylase | FMN | Q96CD2 | COAC | *PPCDC* | |
| Unspecific monooxygenase | Heme-thiolate | Q8N118 | CP4X1 | *CYP4X1* | |
| Phosphatidate phosphatase | Mg(2+) | O14495 | PLPP3 | *PPAP2B* | |
| Phosphatidate phosphatase | Mg(2+) | Q96GM1 | PLPR2 | *PLPPR2* | |
| Apyrase | Ca(2+) | Q5MY95 | ENTP8 | *ENTPD8* | |
| Methylmalonyl-CoA mutase | Cob(II)alamin | P22033 | MUTA | *MUT* | yes |
| Leukotriene-B(4) 20-monooxygenase | Heme-thiolate | P78329 | CP4F2 | *CYP4F2* | |
| Calcium-transporting ATPase | Mg(2+) | Q01814 | AT2B2 | *ATP2B2* | yes |
| Phospholipid-translocating ATPase | Mg(2+) | O43861 | ATP9B | *ATP9B* | |
| 2-(3-amino-3-carboxypropyl)histidine synthase | Iron-sulfur | Q9BZG8 | DPH1 | *DPH1* | |
| Alcohol dehydrogenase | Zn(2+) or Fe cation | P00325 | ADH1B | *ADH1B* | yes |
| Ferroxidase | Cu cation | Q8N4E7 | FTMT | *FTMT* | yes |
| DNA oxidative demethylase | Fe cation | Q6NS38 | ALKB2 | *ALKBH2* | |
| Glycogenin glucosyltransferase | Mn(2+) | O15488 | GLYG2 | *GYG2* | |
| Leukotriene-A(4) hydrolase | Zn(2+) | P09960 | LKHA4 | *LTA4H* | |
| Unspecific monooxygenase | Heme-thiolate | P51589 | CP2J2 | *CYP2J2* | |
| Phospholipid-translocating ATPase | Mg(2+) | P98196 | AT11A | *ATP11A* | |
| Calpain-3 | Ca(2+) | P20807 | CAN3 | *CAPN3* | yes |
| Unspecific monooxygenase | Heme-thiolate | P20815 | CP3A5 | *CYP3A5* | yes |
| Nardilysin | Zn(2+) | O43847 | NRDC | *NRD1* | |
| [Histone H3]-lysine-36 demethylase | Fe(2+) | Q8N371 | KDM8 | *KDM8* | |

| | | | | | |
|---|---|---|---|---|---|
| Carboxypeptidase A | Zn(2+) | P15085 | CBPA1 | *CPA1* | yes |
| Phospholipid-translocating ATPase | Mg(2+) | O43520 | AT8B1 | *ATP8B1* | yes |
| N(1)-acetylpolyamine oxidase | FAD | Q6QHF9 | PAOX | *PAOX* | |
| 4-hydroxyphenylpyruvate dioxygenase | Fe cation | P32754 | HPPD | *HPD* | yes |
| Phospholipid-hydroperoxide glutathione peroxidase | Se(2+) | P36969 | GPX4 | *GPX4* | |
| 3-methyl-2-oxobutanoate dehydrogenase (2-methylpropanoyl-transferring) | Thiamine diphosphate | P12694 | ODBA | *BCKDHA* | yes |
| Unspecific monooxygenase | Heme-thiolate | 87X1C5 | CP2D7 | *0* | |
| Calcium/calmodulin-dependent protein kinase | Ca(2+) | Q6P2M8 | KCC1B | *PNCK* | |
| Phospholipase A(2) | Ca(2+) | P30041 | PRDX6 | *PRDX6* | |
| Superoxide dismutase | Fe cation or Mn(2+) or (Zn(2+) and Cu cation) | P04179 | SODM | *SOD2* | yes |
| Unspecific monooxygenase | Heme-thiolate | P24903 | CP2F1 | *CYP2F1* | |
| Threonine ammonia-lyase | Pyridoxal 5'-phosphate or iron-sulfur | P20132 | SDHL | *SDS* | |
| Phospholipase A(2) | Ca(2+) | O60733 | PLPL9 | *PLA2G6* | yes |
| Cytochrome-b5 reductase | FAD | P00387 | NB5R3 | *CYB5R3* | yes |
| Mitochondrial processing peptidase | Zn(2+) | Q10713 | MPPA | *PMPCA* | |
| Prolyl aminopeptidase | Mn(2+) | P28838 | AMPL | *LAP3* | |
| Galactosylgalactosylxylosylprotein 3-beta-glucuronosyltransferase | Mn(2+) | Q9NPZ5 | B3GA2 | *B3GAT2* | |
| (S)-citramalyl-CoA lyase | Mg(2+) | Q8N0X4 | CLYBL | *CLYBL* | |
| Carbonic anhydrase | Zn(2+) | P23280 | CAH6 | *CA6* | |
| Indoleamine 2,3-dioxygenase | Heme | P14902 | I23O1 | *IDO1* | |
| Phenylalanine 4-monooxygenase | Fe cation | P00439 | PH4H | *PAH* | yes |
| Pyruvate dehydrogenase (acetyl-transferring) | Thiamine diphosphate | P08559 | ODPA | *PDHA1* | yes |
| Unspecific monooxygenase | Heme-thiolate | Q16678 | CP1B1 | *CYP1B1* | yes |
| Methylcrotonoyl-CoA carboxylase | Biotin | Q9HCC0 | MCCB | *MCCC2* | yes |
| Inositol-trisphosphate 3-kinase | Ca(2+) | P27987 | IP3KB | *ITPKB* | |
| Calcium/calmodulin-dependent protein kinase | Ca(2+) | Q13555 | KCC2G | *CAMK2G* | |
| Inositol oxygenase | Fe cation | Q9UGB7 | MIOX | *MIOX* | |
| Methionyl aminopeptidase | Cobalt cation | P50579 | MAP2 | *METAP2* | |
| Glutamate carboxypeptidase II | Zn(2+) | Q04609 | FOLH1 | *FOLH1* | |
| O-phospho-L-seryl-tRNA(Sec):L-selenocysteinyl-tRNA synthase | Pyridoxal 5'-phosphate | Q9HD40 | SPCS | *SEPSECS* | yes |
| Oxoglutarate dehydrogenase (succinyl-transferring) | Thiamine diphosphate | Q96HY7 | DHTK1 | *DHTKD1* | yes |
| Superoxide dismutase | Fe cation or Mn(2+) or (Zn(2+) and Cu cation) | P00441 | SODC | *SOD1* | yes |
| Calcium/calmodulin-dependent protein kinase | Ca(2+) | Q14012 | KCC1A | *CAMK1* | |
| ADAM 17 endopeptidase | Zn(2+) | P78536 | ADA17 | *ADAM17* | yes |
| Stearoyl-CoA 9-desaturase | Fe cation | Q86SK9 | SCD5 | *SCD5* | |
| Flavin-containing monooxygenase | FAD | P31512 | FMO4 | *FMO4* | |
| Urocanate hydratase | NAD(+) | Q96N76 | HUTU | *UROC1* | yes |
| Unspecific monooxygenase | Heme-thiolate | P10632 | CP2C8 | *CYP2C8* | yes |
| FAD-AMP lyase (cyclizing) | Cobalt cation or Mn(2+) | Q3LXA3 | TKFC | *DAK* | |

| | | | | | |
|---|---|---|---|---|---|
| Alcohol dehydrogenase | Zn(2+) or Fe cation | P00326 | ADH1G | *ADH1C* | yes |
| Unspecific monooxygenase | Heme-thiolate | Q9HCS2 | CP4FC | *CYP4F12* | |
| Calcium/calmodulin-dependent protein kinase | Ca(2+) | Q96RR4 | KKCC2 | *CAMKK2* | |
| Alcohol dehydrogenase | Zn(2+) or Fe cation | P11766 | ADHX | *ADH5* | |
| Phospholipase A(2) | Ca(2+) | P0C869 | PA24B | *PLA2G4B* | |
| Hydrogen/potassium-exchanging ATPase | Mg(2+) | P20648 | ATP4A | *ATP4A* | |
| Hydroxymethylbilane synthase | Dipyrromethane | P08397 | HEM3 | *HMBS* | yes |
| Calcium-transporting ATPase | Mg(2+) | Q93084 | AT2A3 | *ATP2A3* | yes |
| Calcium-transporting ATPase | Mg(2+) | O14983 | AT2A1 | *ATP2A1* | yes |
| [Histone H3]-lysine-36 demethylase | Fe(2+) | Q9H6W3 | RIOX1 | *RIOX1* | |
| Arachidonate 5-lipoxygenase | Fe cation | P09917 | LOX5 | *ALOX5* | yes |
| Carbonic anhydrase | Zn(2+) | Q16790 | CAH9 | *CA9* | |
| Glutamate carboxypeptidase II | Zn(2+) | Q9Y3Q0 | NALD2 | *NAALAD2* | |
| Aryldialkylphosphatase | Divalent cation | P27169 | PON1 | *PON1* | yes |
| L-2-hydroxyglutarate dehydrogenase | FAD | Q9H9P8 | L2HDH | *L2HGDH* | yes |
| tRNA(adenine(34)) deaminase | Zn(2+) | Q7Z6V5 | ADAT2 | *ADAT2* | |
| Phospholipase A(2) | Ca(2+) | Q9UP65 | PA24C | *PLA2G4C* | |
| Fructose-bisphosphate aldolase | Zn(2+) | P04075 | ALDOA | *ALDOA* | yes |
| Methionyl aminopeptidase | Cobalt cation | P53582 | MAP11 | *METAP1* | |
| L-methionine (R)-S-oxide reductase | NADPH | Q9Y3D2 | MSRB2 | *MSRB2* | |
| Prostaglandin-I synthase | Heme-thiolate | Q16647 | PTGIS | *PTGIS* | yes |
| Histidine decarboxylase | Pyruvate or pyridoxal 5'-phosphate | P19113 | DCHS | *HDC* | yes |
| 25/26-hydroxycholesterol 7-alpha-hydroxylase | Heme-thiolate | O75881 | CP7B1 | *CYP7B1* | yes |
| Iodotyrosine deiodinase | FMN | Q6PHW0 | IYD1 | *IYD* | yes |
| Adenosylmethionine decarboxylase | Pyruvate | P17707 | DCAM | *AMD1* | |
| [Histone H3]-lysine-36 demethylase | Fe(2+) | Q9UPP1 | PHF8 | *PHF8* | yes |
| Mitochondrial intermediate peptidase | Zn(2+) | Q99797 | MIPEP | *MIPEP* | |
| Pappalysin-1 | Zn(2+) | Q13219 | PAPP1 | *PAPPA* | |
| Pyruvate dehydrogenase (acetyl-transferring) | Thiamine diphosphate | P29803 | ODPAT | *PDHA2* | |
| Phosphoserine transaminase | Pyridoxal 5'-phosphate | Q9Y617 | SERC | *PSAT1* | yes |
| Squalene monooxygenase | FAD | Q14534 | ERG1 | *SQLE* | |
| Beta-carotene 15,15'-dioxygenase | Fe cation | Q9HAY6 | BCDO1 | *BCMO1* | yes |
| Stearoyl-CoA 9-desaturase | Fe cation | O00767 | ACOD | *SCD* | |
| L-dopachrome isomerase | Zn(2+) | P14174 | MIF | *MIF* | yes |
| 3-methyl-2-oxobutanoate dehydrogenase (2-methylpropanoyl-transferring) | Thiamine diphosphate | P21953 | ODBB | *BCKDHB* | yes |
| Fucokinase | Divalent cation | Q8N0W3 | FUK | *FUK* | |
| FAD synthetase | Mg(2+) | Q8NFF5 | FAD1 | *FLAD1* | |
| Cystathionine beta-synthase | Pyridoxal 5'-phosphate | P0DN79 | CBSL | *CBSL* | |
| Glycine hydroxymethyltransferase | Pyridoxal 5'-phosphate | P34897 | GLYM | *SHMT2* | |
| Carboxypeptidase B | Zn(2+) | P15086 | CBPB1 | *CPB1* | |

| | | | | | |
|---|---|---|---|---|---|
| Aconitate hydratase | Iron-sulfur | P21399 | ACOC | *ACO1* | |
| Oxoglutarate dehydrogenase (succinyl-transferring) | Thiamine diphosphate | Q02218 | ODO1 | *OGDH* | |
| Neprilysin | Zn(2+) | Q495T6 | MMEL1 | *MMEL1* | |
| Propionyl-CoA carboxylase | Biotin | P05165 | PCCA | *PCCA* | yes |
| Glutathione peroxidase | Se(2+) | P18283 | GPX2 | *GPX2* | |
| Glutathione peroxidase | Se(2+) | P59796 | GPX6 | *GPX6* | |
| Aryldialkylphosphatase | Divalent cation | Q15166 | PON3 | *PON3* | yes |
| Protein-glutamine gamma-glutamyltransferase | Ca(2+) | P00488 | F13A | *F13A1* | yes |
| 4-aminobutyrate--2-oxoglutarate transaminase | Pyridoxal 5'-phosphate | P80404 | GABT | *ABAT* | yes |
| Carboxypeptidase M | Zn(2+) | P14384 | CBPM | *CPM* | |
| Pyridoxal phosphatase | Mg(2+) | Q8TCD6 | PHOP2 | *PHOSPHO2* | |
| Cytosol alanyl aminopeptidase | Cobalt cation or Zn(2+) | P55786 | PSA | *NPEPPS* | |
| Vitamin D(3) 24-hydroxylase | Heme-thiolate | Q07973 | CP24A | *CYP24A1* | yes |
| Phosphoethanolamine/phosphocholine phosphatase | Mg(2+) or cobalt cation or Mn(2+) | Q8TCT1 | PHOP1 | *PHOSPHO1* | |
| Cholesterol monooxygenase (side-chain-cleaving) | Heme-thiolate | P05108 | CP11A | *CYP11A1* | yes |
| Betaine--homocysteine S-methyltransferase | Zn(2+) | Q93088 | BHMT1 | *BHMT* | |
| Adenosylhomocysteinase | NAD(+) | P23526 | SAHH | *AHCY* | yes |
| Hydrogen/potassium-exchanging ATPase | Mg(2+) | P54707 | AT12A | *ATP12A* | |
| Meprin B | Zn(2+) | Q16820 | MEP1B | *MEP1B* | |
| Methylcytosine dioxygenase | Fe(2+) | Q8NFU7 | TET1 | *TET1* | |
| Cytochrome-c oxidase | Cu cation | P00395 | COX1 | *MT-CO1* | |
| Branched-chain-amino-acid transaminase | Pyridoxal 5'-phosphate | O15382 | BCAT2 | *BCAT2* | |
| Corticosterone 18-monooxygenase | Heme-thiolate | P19099 | C11B2 | *CYP11B2* | yes |
| Hydroperoxy icosatetraenoate isomerase | Fe(2+) | Q9BYJ1 | LOXE3 | *ALOXE3* | yes |
| Lipid-phosphate phosphatase | Mg(2+) | P34913 | HYES | *EPHX2* | yes |
| Meprin A | Zn(2+) | Q16819 | MEP1A | *MEP1A* | |
| 2-aminoadipate transaminase | Pyridoxal 5'-phosphate | Q8N5Z0 | AADAT | *AADAT* | |
| Ferroxidase | Cu cation | P02794 | FRIH | *FTH1* | yes |
| Hyaluronan synthase | Mg(2+) | O00219 | HYAS3 | *HAS3* | |
| Superoxide dismutase | Fe cation or Mn(2+) or (Zn(2+) and Cu cation) | P08294 | SODE | *SOD3* | yes |
| Porphobilinogen synthase | Zn(2+) | P13716 | HEM2 | *ALAD* | yes |
| Phosphopyruvate hydratase | Mg(2+) | A6NNW6 | ENO4 | *ENO4* | |
| Cytosol nonspecific dipeptidase | Zn(2+) | Q96KP4 | CNDP2 | *CNDP2* | |
| Membrane dipeptidase | Zn(2+) | P16444 | DPEP1 | *DPEP1* | |
| Calcium-transporting ATPase | Mg(2+) | P16615 | AT2A2 | *ATP2A2* | yes |
| mRNA N(1)-methyladenine demethylase | Fe(2+) | Q96Q83 | ALKB3 | *ALKBH3* | |
| NAD(P)H dehydrogenase (quinone) | FAD | P15559 | NQO1 | *NQO1* | yes |
| Retinal dehydrogenase | FAD | O94788 | AL1A2 | *ALDH1A2* | yes |
| Ribonucleoside-diphosphate reductase | Fe(3+) or adenosylcob(III)alamin or Mn(2+) | P31350 | RIR2 | *RRM2* | |

| | | | | | |
|---|---|---|---|---|---|
| Calcium/calmodulin-dependent protein kinase | Ca(2+) | Q16566 | KCC4 | *CAMK4* | |
| Sodium/potassium-exchanging ATPase | Mg(2+) | P50993 | AT1A2 | *ATP1A2* | yes |
| Sarcosine oxidase | FAD | Q9P0Z9 | SOX | *PIPOX* | |
| Cholesterol 7-alpha-monooxygenase | Heme-thiolate | P22680 | CP7A1 | *CYP7A1* | yes |
| Peroxidase | Heme | P22079 | PERL | *LPO* | |
| Carbonic anhydrase | Zn(2+) | P00915 | CAH1 | *CA1* | yes |
| Choline dehydrogenase | Pyrroloquinoline quinone | Q8NE62 | CHDH | *CHDH* | |
| Endothelin-converting enzyme 1 | Zn(2+) | P0DPD6 | ECE2 | *ECE2* | |
| Riboflavin kinase | Mg(2+) or Zn(2+) or Mn(2+) | Q969G6 | RIFK | *RFK* | |
| Unspecific monooxygenase | Heme-thiolate | P10635 | CP2D6 | *CYP2D6* | yes |
| Calpain-1 | Ca(2+) | P07384 | CAN1 | *CAPN1* | |
| Phosphatidylserine decarboxylase | Pyruvate or pyridoxal 5'-phosphate | Q9UG56 | PISD | *PISD* | |
| Carbonic anhydrase | Zn(2+) | O43570 | CAH12 | *CA12* | yes |
| Tyrosine 3-monooxygenase | Fe cation | P07101 | TY3H | *TH* | yes |
| [Histone H3]-lysine-36 demethylase | Fe(2+) | Q8NHM5 | KDM2B | *KDM2B* | |
| Cysteine-S-conjugate beta-lyase | Pyridoxal 5'-phosphate | Q6YP21 | KAT3 | *CCBL2* | |
| Cytochrome-b5 reductase | FAD | Q6IPT4 | NB5R5 | *CYB5RL* | |
| Methylcytosine dioxygenase | Fe(2+) | Q6N021 | TET2 | *TET2* | yes |
| Phospholipase A(2) | Ca(2+) | Q68DD2 | PA24F | *PLA2G4F* | |
| Kynureninase | Pyridoxal 5'-phosphate | Q16719 | KYNU | *KYNU* | yes |
| Selenide, water dikinase | Mg(2+) | Q99611 | SPS2 | *SEPHS2* | |
| Serine--pyruvate transaminase | Pyridoxal 5'-phosphate | P21549 | SPYA | *AGXT* | yes |
| ADP-specific glucokinase | Mg(2+) | Q9BRR6 | ADPGK | *ADPGK* | |
| Peroxidase | Heme | A1KZ92 | PXDNL | *PXDNL* | |
| Phospholipid-translocating ATPase | Mg(2+) | P98198 | AT8B2 | *ATP8B2* | |
| Glutamate decarboxylase | Pyridoxal 5'-phosphate | P15104 | GLNA | *GLUL* | yes |
| tRNA 4-demethylwyosine synthase (AdoMet-dependent) | Iron-sulfur | Q9NV66 | TYW1 | *TYW1* | |
| Branched-chain-amino-acid transaminase | Pyridoxal 5'-phosphate | P54687 | BCAT1 | *BCAT1* | |
| Phosphatidate phosphatase | Mg(2+) | Q8NEB5 | PLPP5 | *PPAPDC1B* | |
| Steroid 11-beta-monooxygenase | Heme-thiolate | P15538 | C11B1 | *CYP11B1* | yes |
| Alanine transaminase | Pyridoxal 5'-phosphate | P24298 | ALAT1 | *GPT* | yes |
| D-aminoacyl-tRNA deacylase | Zn(2+) | Q96FN9 | DTD2 | *DTD2* | |
| dTDP-glucose 4,6-dehydratase | NAD(+) | O95455 | TGDS | *TGDS* | |
| Protein-glutamine gamma-glutamyltransferase | Ca(2+) | P21980 | TGM2 | *TGM2* | yes |
| Isovaleryl-CoA dehydrogenase | FAD | P26440 | IVD | *IVD* | yes |
| Molybdenum cofactor sulfurtransferase | Pyridoxal 5'-phosphate | Q96EN8 | MOCOS | *MOCOS* | yes |
| Glutaryl-CoA dehydrogenase (ETF) | FAD | Q92947 | GCDH | *GCDH* | yes |
| Serine C-palmitoyltransferase | Pyridoxal 5'-phosphate | O15270 | SPTC2 | *SPTLC2* | yes |
| 1,8-cineole 2-exo-monooxygenase | Heme-thiolate | P08684 | CP3A4 | *CYP3A4* | yes |
| Phospholipid-translocating ATPase | Mg(2+) | O75110 | ATP9A | *ATP9A* | |

| | | | | | |
|---|---|---|---|---|---|
| Sphinganine-1-phosphate aldolase | Pyridoxal 5'-phosphate | O95470 | SGPL1 | *SGPL1* | |
| 2-methyl-branched-chain-enoyl-CoA reductase | FAD | P45954 | ACDSB | *ACADSB* | yes |
| Renalase | FAD | Q5VYX0 | RNLS | *RNLS* | yes |
| Sodium/potassium-exchanging ATPase | Mg(2+) | P13637 | AT1A3 | *ATP1A3* | yes |
| Unspecific monooxygenase | Heme-thiolate | Q16696 | CP2AD | *CYP2A13* | |
| Phosphopyruvate hydratase | Mg(2+) | P09104 | ENOG | *ENO2* | |
| Glycine hydroxymethyltransferase | Pyridoxal 5'-phosphate | P34896 | GLYC | *SHMT1* | |
| Prostaglandin-E synthase | Glutathione | Q9H7Z7 | PGES2 | *PTGES2* | |
| Carbonic anhydrase | Zn(2+) | Q9ULX7 | CAH14 | *CA14* | |
| Carbonic anhydrase | Zn(2+) | P07451 | CAH3 | *CA3* | |
| Acyl-CoA oxidase | FAD | Q15067 | ACOX1 | *ACOX1* | yes |
| Phosphatidate phosphatase | Mg(2+) | O43688 | PLPP2 | *PPAP2C* | |
| Calcium/calmodulin-dependent protein kinase | Ca(2+) | Q96NX5 | KCC1G | *CAMK1G* | |
| Cholate--CoA ligase | Mg(2+) | Q9Y2P5 | S27A5 | *SLC27A5* | yes |
| Metallocarboxypeptidase D | Zn(2+) | O75976 | CBPD | *CPD* | |
| DNA N(6)-methyladenine demethylase | Fe(3+) | Q13686 | ALKB1 | *ALKBH1* | |
| Phosphatidate phosphatase | Mg(2+) | O14494 | PLPP1 | *PPAP2A* | |
| Phosphatidylinositol-3,4,5-trisphosphate 3-phosphatase | Mg(2+) | Q6XPS3 | TPTE2 | *TPTE2* | |
| 3-mercaptopyruvate sulfurtransferase | Zn(2+) | P25325 | THTM | *MPST* | |
| Spermine oxidase | FAD | Q9NWM0 | SMOX | *SMOX* | |
| Cystathionine gamma-lyase | Pyridoxal 5'-phosphate | P32929 | CGL | *CTH* | yes |
| Galactosylxylosylprotein 3-beta-galactosyltransferase | Mn(2+) | Q96L58 | B3GT6 | *B3GALT6* | yes |
| Glutathione peroxidase | Se(2+) | O75715 | GPX5 | *GPX5* | |
| Methylmalonyl-CoA decarboxylase | Biotin | Q9NTX5 | ECHD1 | *ECHDC1* | |
| Phospholipid-translocating ATPase | Mg(2+) | O60423 | AT8B3 | *ATP8B3* | |
| Arachidonate 12-lipoxygenase | Fe cation | P18054 | LOX12 | *ALOX12* | |
| L-methionine (R)-S-oxide reductase | NADPH | Q8IXL7 | MSRB3 | *MSRB3* | yes |
| D-lactate dehydrogenase (cytochrome) | FAD | Q86WU2 | LDHD | *LDHD* | |
| Thimet oligopeptidase | Zn(2+) | P52888 | THOP1 | *THOP1* | |
| Protoporphyrinogen oxidase | FAD | P50336 | PPOX | *PPOX* | yes |
| Unspecific monooxygenase | Heme-thiolate | P13584 | CP4B1 | *CYP4B1* | |
| 4-nitrophenol 2-hydroxylase | Heme-thiolate | P05181 | CP2E1 | *CYP2E1* | yes |
| Quercetin 2,3-dioxygenase | Fe cation or Cu cation | O00625 | PIR | *PIR* | |
| GDP-mannose 4,6-dehydratase | NAD(+) | O60547 | GMDS | *GMDS* | |
| Apyrase | Ca(2+) | P49961 | ENTP1 | *ENTPD1* | yes |
| Galactosylgalactosylxylosylprotein 3-beta-glucuronosyltransferase | Mn(2+) | O94766 | B3GA3 | *B3GAT3* | yes |
| Phosphatidate phosphatase | Mg(2+) | Q6T4P5 | PLPR3 | *PLPPR3* | |
| Peptide-aspartate beta-dioxygenase | Fe(2+) | Q12797 | ASPH | *ASPH* | |
| Unspecific monooxygenase | Heme-thiolate | P98187 | CP4F8 | *CYP4F8* | |
| Adenosylhomocysteinase | NAD(+) | Q96HN2 | SAHH3 | *AHCYL2* | |

| | | | | | |
|---|---|---|---|---|---|
| Unspecific monooxygenase | Heme-thiolate | P24462 | CP3A7 | *CYP3A7* | yes |
| Xaa-Pro dipeptidase | Mn(2+) | P12955 | PEPD | *PEPD* | yes |
| UDP-glucose 4-epimerase | NAD(+) | Q14376 | GALE | *GALE* | yes |
| Calcium-transporting ATPase | Mg(2+) | P20020 | AT2B1 | *ATP2B1* | |
| Ferroxidase | Cu cation | Q16595 | FRDA | *FXN* | yes |
| Aminopeptidase B | Zn(2+) | Q9H4A4 | AMPB | *RNPEP* | |
| 5-phosphonooxy-L-lysine phospho-lyase | Pyridoxal 5'-phosphate | Q8IUZ5 | AT2L2 | *PHYKPL* | |
| Cystathionine beta-synthase | Pyridoxal 5'-phosphate | P35520 | CBS | *CBS* | yes |
| Glutathione peroxidase | Se(2+) | P07203 | GPX1 | *GPX1* | yes |
| Kynurenine 3-monooxygenase | FAD | O15229 | KMO | *KMO* | |
| Carbonic anhydrase | Zn(2+) | P00918 | CAH2 | *CA2* | yes |
| (R)-limonene 6-monooxygenase | Heme-thiolate | P11712 | CP2C9 | *CYP2C9* | yes |
| Phospholipase A(2) | Ca(2+) | P14555 | PA2GA | *PLA2G2A* | yes |
| Glutathione peroxidase | Se(2+) | Q96SL4 | GPX7 | *GPX7* | |
| Phospholipase A(2) | Ca(2+) | Q9HCN3 | TMM8A | *TMEM8A* | |
| Glutathione peroxidase | Se(2+) | P22352 | GPX3 | *GPX3* | |
| Carboxypeptidase A2 | Zn(2+) | P48052 | CBPA2 | *CPA2* | |
| L-methionine (R)-S-oxide reductase | NADPH | Q9NZV6 | MSRB1 | *MSRB1* | |
| Cysteine transaminase | Pyridoxal 5'-phosphate | P17174 | AATC | *GOT1* | yes |
| S2P endopeptidase | Zn(2+) | O43462 | MBTP2 | *MBTPS2* | yes |
| Lipoyl synthase | Iron-sulfur | O43766 | LIAS | *LIAS* | yes |
| tRNA (N(6)-L-threonylcarbamoyladenosine(37)-C(2))-methylthiotransferase | Iron-sulfur | Q5VV42 | CDKAL | *CDKAL1* | yes |
| Peroxidase | Heme | P11678 | PERE | *EPX* | yes |
| Phosphatidate phosphatase | Mg(2+) | Q7Z2D5 | PLPR4 | *PLPPR4* | |
